# A proteomic signature of oocyte quality from models of varying oocyte developmental competence

**DOI:** 10.64898/2026.01.20.700687

**Authors:** Emily R. Frost, Dulama Richani, Anne Poljak, Ananya Vuyyuru, Xuihua Liao, Elise Georgiou, J M Binuri Gunasekara, Bettina P. Mihalas, Irene E. Sucquart, Kaushiki Kadam, Lindsay E. Wu, Robert B. Gilchrist

**Affiliations:** Fertility & Research Centre, Discipline of Women’s Health, School of Clinical Medicine, University of New South Wales Sydney, Australia; Bioanalytical Mass Spectrometry Facility, Mark Wainwright Analytical Centre, University of New South Wales Sydney, Australia; School of Biomedical Sciences, University of New South Wales Sydney, Australia

**Keywords:** Capacitation-IVM (CAPA-IVM), *in vitro* maturation (IVM), folliculogenesis, assisted reproductive technologies, proteomics

## Abstract

**Study question:** Which proteins underpin oocyte developmental competence, as modelled by oocytes of variable competence matured *in vivo*, or matured *in vitro* under different conditions (capacitation *in vitro* maturation (CAPA) or standard *in vitro* maturation (IVM))?

**Summary answer:** Significant differences in the global proteome were observed in both oocytes and their corresponding cumulus cells depending on the mode of oocyte maturation, with key variations in eukaryotic translation, autophagy and endocytosis pathways within oocytes, and changes in reactive oxygen species detoxification and serine biosynthesis in cumulus cells.

**What is known already:** Within the ovarian follicle, mammalian oocytes must acquire the necessary molecular machinery to support successful fertilisation and embryonic development. Close contact with the surrounding cumulus cells ensures coordinated nuclear and cytoplasmic maturation of the oocyte, along with the accumulation of proteins stored within the oocyte in cytoplasmic lattices and endo-lysosomal vesicular assemblies.

**Study design, size, duration:** This basic science study utilised a mouse model to assess proteomic changes across three oocyte competence models. Key proteins identified in mouse oocytes were also assessed in discarded immature human germinal vesicle (GV) oocytes and MII oocytes following rescue-IVM. Three oocyte maturation methods were tested: i) *in vivo* maturation, (ii) CAPA and (iii) standard IVM. *In vivo* maturation served as a positive control group, whereby metaphase II (MII) mature oocytes were collected from mice stimulated with pregnant mare serum gonadotropin (PMSG) and triggered with human chorionic gonadotropin (hCG), simulating full ovarian stimulation. For the *in vitro* maturation groups, immature cumulus oocyte complexes (COCs) were collected from mildly stimulated (23 hr PMSG) mice. For the standard IVM group, immature COCs were matured in media containing amphiregulin and epiregulin for 18 hours. For the CAPA group, COCs were held for 24 hours in pre-IVM conditions in the presence of c-type natriuretic peptide (CNP), oestradiol, insulin and follicle stimulating hormone (FSH), and then matured via IVM in media containing FSH, amphiregulin and epiregulin. Four biological replicates were performed for mouse proteomics experiments, three biological replicates performed for mouse immunocytochemistry experiments and six replicates were performed for embryology experiments.

**Participants/materials, settings, methods:** Four to six-week-old C57BL/6JAusb mice were used for all mouse experiments. Embryology outcomes were used to confirm the variation in oocyte developmental competence between the three maturation groups. For the *in vivo*, CAPA and IVM groups, mature MII COCs were collected and separated into oocytes and cumulus cells. Oocytes and cumulus cells were subjected to mass spectrometry and bioinformatic analysis was performed using Proteome Discoverer and Ingenuity Pathway Analysis, with data validated by immunofluorescence. To assess conservation of proteins in human oocytes, 49 oocytes were collected from 36 patients following assisted reproduction technology (ART) cycles and subject to immunofluorescence. Rescue-IVM was also performed with half of the human oocyte cohort to obtain MII oocytes.

**Main results and the role of chance:** Proteomic profiling identified around 1600 proteins in mouse oocytes and 3100 in mouse cumulus cells across all three treatment groups (at least 2 peptides per protein). Differential expression analysis and pattern analysis collectively revealed a signature of proteins that were consistently differentially expressed between *in vivo* and *in vitro* oocyte maturation systems (log2FC of ± 1 and a p-value ≤ 0.05). These subsets of proteins were mapped to biological processes including eukaryotic translation, autophagy and endocytosis pathways within oocytes. Orthogonal validation of clathrin, ribosomal protein L24 and eukaryotic initiation factor 2A supported the proteomic findings and expression was conserved in human oocytes. Changes in reactive oxygen species detoxification and serine biosynthesis were observed in mouse cumulus cells, with fluorescence intensity changes in ferredoxin-1 and phosphoglycerate dehydrogenase supporting the dysregulation of cumulus cell processes during *in vitro* maturation.

**Large scale data:** The mass spectrometry data are available via ProteomeXchange with identifier PXD073269.

**Limitations, reasons for caution:** The foundational mechanisms of oocyte developmental competence remain elusive, particularly in humans where MII oocytes are heterogenous in quality within the same stimulation cycle and patient. In this study, C57Bl6/J mice were used as the model species, allowing precise control over differing models of oocyte quality and capacity to analyse large numbers of oocytes. However, care is required when interpreting the significance of these findings in mice to mechanisms regulating human oocyte quality. Nonetheless, the *in vivo* stimulation and both IVM protocols used in this study are clinically relevant and developmentally matched. This study has also not addressed oocyte developmental competence in gonadotropin-free IVM oocytes, which is now a clinical reality.

**Wider implications of the findings:** This study confirms that mouse oocytes, matured *in vitro* in two clinically relevant systems, show reduced developmental competence when compared to *in vivo* matured oocytes. Through examination of the global proteome in oocytes, molecular pathways including eukaryotic translation, autophagy and endocytosis were dysregulated in *in vitro* oocytes. Recent findings have revealed the critical role of these pathways to developmental competence in the context of *in vivo* development. In cumulus cells, changes in reactive oxygen species detoxification and serine biosynthesis were observed, adding to the extensive knowledge around metabolic activity in cumulus cells as a critical facet of oocyte quality. Combined, this data suggests that the necessary processes of protein storage and degradation in oocytes and metabolism in cumulus cells constitute important components of oocyte quality. These processes appear suboptimal in current IVM systems, providing a future research direction to optimise IVM protocols with consideration to these protein pathways.

**Study funding/competing interests:** This study was funded by a National Health and Medical Research Council Investigator Fellowship (APP1023210) awarded to R.B.G. and by a gift from Open Philanthropy. The following competing interests are declared: R.B.G.is a consultant to Dioseve Inc.. L.E.W is a co-founder, shareholder, director and advisor of Jumpstart Fertility Inc.. L.E.W. is also an advisor and shareholder in EdenRoc Sciences, the parent company of Metro Biotech NSW and Metro Biotech, and in Life Biosciences LLC and its daughter companies. His UNSW Industry Scientia position is partly funded by Proto Axiom. All other authors have no competing interests to disclose.

## Introduction

Over many decades, assisted reproductive technology (ART) treatment strategies has been refined to improve the efficiency of protocols and importantly reduce the time to live birth for patients. Despite this, the global pregnancy rate per aspiration for *in vitro* fertilisation (IVF)/ intracytoplasmic sperm injection (ICSI) cycles remains around 26% (Chambers, et al., 2021). A principal determinant of ART success is the retrieval of a population of high-quality oocytes that can undergo fertilisation and support embryo and fetal development. Despite extensive research effort, it is still unclear what key molecular events constitute a high-quality oocyte. Thus, there are no therapies to improve oocyte quality and clinical assessment of oocytes remains primarily restricted to morphological assessments (Lemseffer, et al., 2022, Sciorio, et al., 2022). Elucidating a molecular signature of oocyte developmental competence will provide opportunities to further improve ART outcomes and diagnostic methods to quantify gamete potential.

The capacity of an oocyte to undergo fertilisation and support embryo development is referred to as oocyte developmental competence. Developmental competence covers a broad range of contributions to oocyte quality. This includes the attainment of all molecular machinery necessary to support the oocyte through meiosis and early embryo development, up to and beyond activation of the embryonic genome (Tan and Dunning, 2022). Oocyte developmental competence is initiated early in folliculogenesis and is continually acquired as the oocyte stores maternal transcripts, proteins and metabolites before entering a period of transcriptional silence in late folliculogenesis (Li, et al., 2010, Zhang, et al., 2018). It is widely accepted that several cellular and molecular processes are critical foundations for a healthy oocyte, including; bidirectional oocyte/cumulus cell communication, accumulation of mitochondria and appropriate metabolic activity, accumulation of proteins and RNAs, capacity to elicit appropriate damage responses, resumption of the meiotic cell cycle, proper chromosome condensation, spindle assembly and faithful chromosome segregation (reviewed in (Conti and Franciosi, 2018)). Recent studies have highlighted that efficient removal of unwanted maternal products is also a key aspect of oocyte developmental competence (Sha, et al., 2020, Zaffagnini, et al., 2024, Zaffagnini, et al., 2025). Importantly, all these processes must be precisely controlled to ensure coordinated nuclear and cytoplasmic maturation, and this heavily relies on signalling from the associated cumulus cells (Richani, et al., 2021). Cumulus cells play a vital role in supplying metabolites, nutrients and other paracrine signalling molecules to the oocyte, with disruptions in communication and function resulting in poor quality oocytes and decreased rates of embryo formation and development (Dvoran, et al., 2024, Fatehi, et al., 2002, Geshi, et al., 2000, Hashimoto, et al., 1998, Richani, et al., 2019, Wang, et al., 2025). Despite an ever-expanding knowledge base of the components of oocyte and cumulus cell quality, a molecular signature of a high-quality oocyte remains elusive and other undiscovered processes are likely contributors.

Investigating the underlying mechanisms of human developmental competence is a technical challenge, hence embryo development and pregnancy outcomes remain the gold standard measure for oocyte quality. In humans, interventions to improve oocyte development cannot be readily assessed for developmental competence due to ethical concerns in generating human embryos for research. However, substantial differences in developmental competence exist between oocyte maturation strategies used in clinical ART (Vuong, et al., 2020a, Vuong, et al., 2020b). Collecting human oocytes from antral follicles and maturing them using *in vitro* maturation (IVM) prior to IVF is a low-intervention approach. Despite successful IVM implementation in the cattle ART industry, human IVM has seen slow clinical progress, primarily attributed to a lower number of resultant embryos and lower cumulative birth rate compared with oocytes from stimulated IVF patients (Walls, et al., 2014). A novel biphasic approach to IVM, termed CAPA-IVM, where the oocyte is ‘capacitated’ using a pre-IVM culture step to gain developmental competence *in vitro* before IVM, is an improvement on standard IVM and has been implemented successfully in clinical practice (Sanchez, et al., 2019, Sánchez, et al., 2017, Vuong, et al., 2025, Vuong, et al., 2020b). While a suitable alternative for women with high antral follicle count or at risk of ovarian hyperstimulation syndrome, cumulative birth rate remains lower for CAPA-IVM patients compared with traditional IVF patients (Gilchrist, et al., 2024, Vuong, et al., 2020a, Vuong, et al., 2025, Vuong, et al., 2020b). These clinical differences in grades of human oocyte developmental competence can be mimicked in a mouse model, where studies have utilised IVM culture systems to investigate the transcriptomic contributions to developmental competence (Dvoran, et al., 2024, Wang, et al., 2025).

Here, we interrogate constituents of oocyte developmental competence by comparing the proteomic signature of mature mouse metaphase II (MII) cumulus-oocyte complexes (COCs) of differing intrinsic qualities, as matured by various maturation protocols. *In vivo* ovulated MII COCs (termed *in vivo* throughout the manuscript), *in vitro* matured MII COCs (IVM) and MII COCs *in vitro* matured using the CAPA-IVM biphasic IVM system (CAPA) were the chosen protocols, as they are used clinically and importantly display varying levels of developmental competence (Sánchez, et al., 2017, Vuong, et al., 2020b). We employed label-free mass spectrometry to analyse the proteomes of mouse oocytes and cumulus cells separately. Proteomic profiling identified novel proteins in key biological pathways that are altered between *in vivo* and *in vitro* maturation strategies, revealing a protein signature of oocyte developmental competence. Immunofluorescence staining validated components of the proteomic signature and confirmed that these key targets are also conserved in human oocytes.

## Experimental Procedures

### Human oocyte collection

Approval of this study was granted by the South Eastern Sydney Local Health District Ethics Committee (2021-ETH00018). Informed consent for the donation of discarded oocytes was provided by couples undergoing ICSI at the Fertility & Research Centre in the Royal Hospital for Women, Sydney. Ovarian stimulation was performed with FSH using either gonadotropin-releasing hormone (GnRH) agonist or antagonist protocols, according to standard clinical procedures (Figure 1B). Following COC collection, denudation and staging, immature oocytes were identified and labelled as research material. Only immature oocytes that could not be used for treatment and would have otherwise been disposed of were used for this study. Cells were transported at 37°C in a portable incubator for approximately 15 mins to the research laboratory and staged as either germinal vesicle (GV) or metaphase-I (MI). A total of 49 oocytes collected from 36 patients, between June 2025 to November 2025, were used in this study.

**Figure 1.**
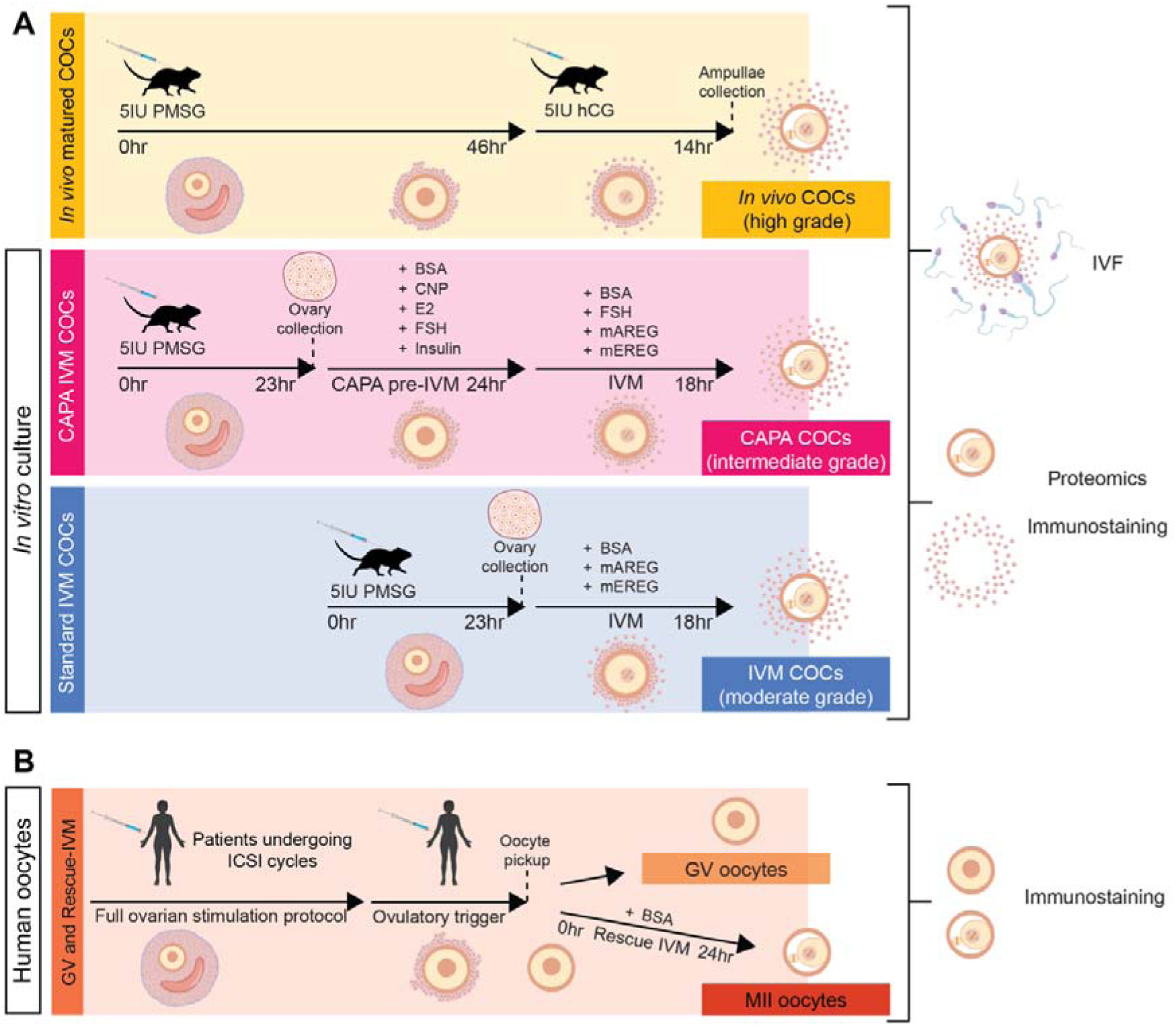
Schematic showing experimental treatment schedules, ovary collection timepoints and endpoints assessed. A) For *in vivo* maturation (yellow box), 4 to 6-week-old female C57Bl/6J mice were stimulated with pregnant mare serum gonadotropin (PMSG) for 46 hours, triggered with human chorionic gonadotropin (hCG) and mature cumulus-oocyte complexes (COCs) were retrieved 14 hours post-trigger. For *in vitro* culture groups, 4 to 6-week-old female C57Bl/6J mice were stimulated with PMSG and ovaries collected 23 hours later to retrieve immature COCs. For the CAPA treatment group (pink box) COCs were cultured in CAPA pre-medium for 24 hours and then *in vitro* maturation (IVM) medium containing listed additives for 18 hours. For the standard IVM treatment group (blue box), COCs were cultured in IVM medium for 18 hours. COCs were then used intact for embryology assessments or separated into oocytes and cumulus cells for proteomics and validation experiments. B) For protein assessment in human oocytes, patients underwent a full stimulation protocol for an intracytoplasmic sperm injection (ICSI) cycle as per standard clinical procedures. Following oocyte pickup, immature germinal vesicle (GV) oocytes were collected from the clinic and transported to the lab. Oocytes were allocated into two groups, either fixed at the GV stage, or oocytes underwent rescue-IVM for 24hrs to reach the metaphase II (MII) stage. Immunostaining experiments were performed on both cohorts of oocytes.

### Human oocyte maturation

To obtain mature MII human oocytes, GV oocytes were subjected to rescue-IVM (Figure 1B). GV oocytes were washed 3 times with standard IVM media containing bicarbonate-buffered alpha minimum essential medium (αMEM) (catalogue 12000063, Gibco, with custom formulation to replace L-glutamine with L-alanyl-L-glutamine at 2 mM concentration) + 3mg/mL fatty-acid free bovine serum albumin (BSA, catalogue A8806, Sigma-Aldrich, Merck, Darmstadt, Germany) supplemented with 50 μg/mL gentamycin (catalogue G1914, Sigma-Aldrich). Oocytes were then transferred to pre-equilibrated standard IVM medium in 4-well culture dishes (NUNC, Thermo Scientific, Roskilde, Denmark) and cultured for 24 hours in 37°C, 5% CO_2_ in humidified air. Cells were then prepared for immunocytochemistry as detailed below.

### Animals

C57BL/6JAusb female mice were used for all experiments and supplied by Australian BioResources, Mossvale, NSW, Australia. All experiments involving the use of animals were conducted in accordance with the Australian Code of Practice for the Care and Use of Animals for Scientific Purposes. Mice were sustained with a 12-hour light-dark cycle, had ad libitum access to food and water and were monitored weekly. All procedures were approved by the UNSW Animal Care & Ethics Committee (ethics numbers 17/105A, 22/40A and 6531).

### Mouse experimental model and cumulus-oocyte complex (COC) collection

This study employed three models of mouse oocyte quality, or oocyte developmental competence (Figure 1A): 1) high grade oocytes matured *in vivo* following superovulation, 2) intermediate grade oocytes matured *in vitro* using CAPA-IVM, and 3) moderate grade oocytes matured *in vitro* using standard IVM. For all treatment groups, four to six-week-old C57BL/6JAusb female mice were stimulated with 5 IU pregnant mare serum gonadotrophin (PMSG, Folligon; Intervet, Boxmeer, The Netherlands and PMSG; Prospec-Tany Technogene Ltd, Israel). For CAPA-IVM and standard IVM treatment group collections, ovaries were harvested 23 hours after PMSG injection (Figure 1A) and placed in HEPES-buffered αMEM (catalogue 12000063, Gibco, with custom formulation to replace L-glutamine with L-alanyl-L-glutamine at 2 mM concentration) containing 3 mg/mL BSA (catalogue A7906, Sigma-Aldrich), 50 μg/mL gentamycin (catalogue G1914, Sigma-Aldrich) and 100 µM 3-Isobutyl-1- methylxanthine (IBMX, catalogue I5879, Sigma-Aldrich). Ovarian preovulatory follicles were punctured with a 1mL insulin syringe attached to a 29-gauge needle to release COCs from the ovary and COCs were gently pipetted into fresh αMEM media + 3 mg/mL BSA + 100 µM 3-Isobutyl-1-methylxanthine + 50 μg/mL gentamycin. For *in vivo* matured oocyte collections (termed *in vivo* thereafter), at 46h post-PMSG injection, mice were injected with 5 IU human chorionic gonadotrophin (hCG; Chorulon, MSD Animal Health, Australia) to induce ovulation. Ovulated COCs were collected from the oviductal ampullae using a 1mL insulin syringe attached to a 29-gauge needle and placed into αMEM + 3 mg/mL BSA + 50 μg/mL gentamycin media. To remove cumulus cells from ovulated oocytes, COCs were briefly incubated for 1 min in αMEM medium supplemented with 3 mg/mL BSA and hyaluronidase (catalogue H4272, Sigma-Aldrich).

### Mouse oocyte IVM and pre-IVM

#### CAPA pre-IVM and IVM system

Following COC collection for the CAPA-IVM treatment group (Figure 1A), COCs were washed 3 times in HEPES-buffered αMEM + 3 mg/mL BSA media to remove the meiotic maturation inhibitor IBMX. CAPA pre-IVM media was prepared with bicarbonate-buffered αMEM + 3 mg/mL BSA supplemented with 50 μg/mL gentamycin (catalogue G1914, Sigma-Aldrich), 2.5 mIU/mL recombinant human FSH (Follitropin Beta; Puregon, Organon, Oss, the Netherlands), 5 ng/mL insulin (catalogue I9278, Sigma-Aldrich), 10 nM estradiol (E2; catalogue E4389, Sigma-Aldrich), and 25 nM c-type natriuretic peptide (CNP, catalogue #3520, Tocris Bioscience, Abingdon, UK). For experiments where *in vitro* embryo production was the endpoint, 1 mg/mL fetuin (catalogue F3385, Sigma-Aldrich) was also added to the media to prevent zona pellucida hardening. COCs were then washed 3 times in the CAPA pre-IVM media, before they were transferred to 500 μL pre-equilibrated CAPA pre-IVM medium (up to 50 COCs per 500 μL of media) in 4-well culture dishes (NUNC, Thermo Scientific) for 24 hours in 37°C, 5% CO_2_ in humidified air. Following CAPA pre-IVM, COCs were washed 3 times in CAPA IVM media (bicarbonate-buffered αMEM + 3mg/mL fatty-acid free BSA supplemented with 50 μg/mL gentamycin, 2.5 mIU/mL FSH, 50 ng/mL recombinant mouse amphiregulin (mAREG; catalogue #989-AR, R&D Systems) and 50 ng/mL recombinant epiregulin (mEREG; catalogue #1068-ER, R&D Systems)). For experiments where *in vitro* embryo production was the endpoint, 1 mg/mL fetuin was also added to the IVM media. COCs were then cultured in 500 μL CAPA IVM medium as described earlier for 18 hours in 37°C, 5% CO_2_ in humidified air. Following CAPA-IVM, COCs were either prepared for IVF, immunocytochemistry or COCs were denuded using a narrow glass pipette, providing separate oocyte and cumulus cell samples. Cells were either processed for immunocytochemistry or snap frozen for proteomics (see below). Figure 1A shows an illustration of the various treatment groups, their culture components and culture timing.

### Standard IVM system

Following COC collection for the IVM treatment group (Figure 1A), COCs were washed 3 times in HEPES-buffered αMEM + 3 mg/mL BSA media. COCs were then washed 3 times in standard IVM media, containing bicarbonate-buffered αMEM + 3mg/mL BSA supplemented with 50 μg/mL gentamycin, 50 ng/mL mAREG and 50 ng/mL mEREG. For experiments where *in vitro* embryo production was the endpoint, 1 mg/mL fetuin was also added. COCs were then cultured in 500 μL standard IVM medium for 18 hours in 37°C, 5% CO_2_ in humidified air. Following IVM, COCs were treated as per CAPA-IVM COCs, as described above (Figure 1A).

### Mouse *in vitro* embryo production

Performing IVF of COCs after CAPA-IVM, standard IVM or *in vivo* maturation was used as an assessment of oocyte quality, as it provides a measure of preimplantation embryo development. Following CAPA-IVM, standard IVM or *in vivo* maturation, COCs were fertilised and the resulting embryos were cultured for six days as previously described (Stocker, et al., 2020). All media (Research Wash, Research Fert and Research Cleave) were purchased from ART Lab Solutions (Adelaide, Australia). Briefly, COCs were washed 3 times in Research Wash media, supplemented with 4 mg/mL fatty-acid free BSA, then washed 3 times in Research Fert media, supplemented with 4 mg/mL fatty-acid free BSA. Sperm used to fertilise oocytes was collected fresh from hybrid CBB6F1 males > 12 weeks of age and capacitated in Research Fert + BSA for at least one hour. COCs were co-incubated with capacitated sperm for four hours at 37°C with 5% O_2_, 6% CO_2_ and 89% N_2_ in Research Fert media + BSA. Presumptive zygotes were washed 3 times in Research Wash media + BSA, then washed 3 times in Research Cleave media, supplemented with 4 mg/mL fatty-acid free BSA. Embryos were cultured in Research Cleave media + BSA covered by a layer of washed paraffin oil, at a density of one embryo per 2 μL media. Fertilisation and blastocyst formation rates were assessed at days 2, 5 and 6 post-fertilisation.

### Protein extraction and mass spectrometry for proteomic analysis

Four biological replicates were included in the proteomic analysis for each treatment group (*in vivo*, CAPA-IVM and IVM). Oocytes and cumulus cells were analysed separately. The number of COCs used for each replicate is listed in Supplemental Table 1. Mass spectrometry methodology and bioinformatics analysis was performed as previously described (Richani, et al., 2024). Briefly, cells were lysed and protein extracted in RIPA buffer containing protease inhibitors; 0.1 mM phenylmethylsulfonyl fluoride (PMSF) and 10 mM ethylenediaminetetraacetic acid (EDTA). Samples were run on a QExactive mass spectrometer (Thermo Electron, Bremen, Germany), with peptide separation on a Dionex UltiMate 3000 nano-liquid chromatography (nano-LC) system (ThermoScientific, Waltham, MA), equipped with an autosampler (Dionex, Amsterdam, Netherlands).

### Bioinformatics analysis

Proteome Discoverer 2.4 (Thermo Fisher Scientific, Waltham, MA) was used to perform database searching of all mass spectrometry raw files for identifying peptides/proteins and quantifying peak abundances (areas) for the purpose of determining protein expression change across treatments. For the processing workflow, the following parameters were used: retention time alignment was performed with a maximum 3 min retention time shift and a minimum signal:noise threshold of 5, use of 3 search engines (Mascot, Sequest and MS Amanda), protein level false discovery rate (FDR) was reported based on a concatenated decoy database (FDR set between 1 – 5 %), with an identification requirement of a minimum of 2 peptide sequences per protein. For the consensus workflow, the following parameters were used: precursor quantification based on peak area and use of unique (but not razor) peptides, normalization was based on total peptide amount and protein abundance calculated as summed abundances, and relative quantification was presented as pairwise peak abundance (area) ratios of treatment groups compared to each other. No data imputation was used, and a t-test (background based) was used to determine expression differences. Principal component analysis and heatmaps were generated within the Proteome Discoverer 2.4 software. The protein list was exported from Proteome Discoverer 2.4 and refined to include only proteins with a quantitative value in at least 3 replicates within a treatment group and a minimum of two unique peptides. From this refined list of proteins, differentially expressed proteins were determined by pairwise comparison, with at least a log2FC of ± 1 (i.e., minimum 2-fold expression difference) and a p-value ≤ 0.05. Volcano plots of differentially expressed proteins were generated in Excel. The lists of differentially expressed proteins were further interrogated using Ingenuity Pathway Analysis software (Qiagen, Hilden, Germany) to identify canonical pathways and upstream regulators associated with the differentially expressed proteins (Krämer, et al., 2014).

### Immunocytochemistry

Mouse COCs and human oocytes were prepared for immunostaining as previously described with minor modifications (Mihalas, et al., 2024). Briefly, COCs and oocytes were quickly washed in PHEM buffer (60mM free acid PIPES, 25mM free acid HEPES, 10mM EGTA and 4mM magnesium sulphate heptahydrate; pH 6.9) containing 0.25% Triton X-100 at 37°C before proceeding with the immunocytochemistry protocol. Following the wash, cells were fixed with 4% formaldehyde in PHEM buffer with 0.25% Triton X-100 for 30 min. Cells were then permeabilised in phosphate-buffered saline with 0.25% Triton X-100 (PBST) for 15 min at room temperature, before blocking with 3% BSA in PBST for 2 hours at room temperature. Cells were incubated in primary antibodies diluted in 3% BSA and 7% goat serum in PBST overnight at 4°C. Antibody details and dilutions are listed in Supplemental Table 2. Following overnight incubation, cells were rinsed 2 times in 1% BSA in PBST before a 1 hr wash in 1% BSA in PBST, before incubation with species-matched secondary antibodies (goat anti-rabbit Alexa Fluor 647 (1:500; A-21244, ThermoFisher Scientific) and goat anti-rabbit Alexa Fluor 488 (1:500; A-11008, ThermoFisher Scientific). Secondary antibodies were diluted in 3% BSA and 7% goat serum in PBST and incubated for 2 hours at room temperature. Following secondary antibody staining, COCs were co-stained with Alexa Fluor 488 Phalloidin (A12379, ThermoFisher Scientific) in 1% BSA and 7% donkey serum or 7% goat serum diluted in PBST overnight at 4°C. Oocytes and COCs were counterstained with the nuclear stain Hoechst (1:2000, H3570, ThermoFisher Scientific) and mounted on FluoroDishes (FD35-100, WPI) in 0.4% polyvinyl alcohol (PVA) in PBS under paraffin oil.

### Confocal microscopy and image analysis

Oocytes and COCs were imaged using the LSM980 laser scanning confocal microscope (Zeiss). Images were taken using the 40x oil objective (Zeiss). To prevent emission overlap, images were acquired sequentially using separate, wavelength-sensitive matched detectors and fluorophores with minimal emission spectra overlap were used. Any potential crosstalk was corrected by adjusting the emission spectra wavelengths and sampling parameters satisfied the Nyquist criterion. For oocytes and COCs, single-slice images were taken across the equatorial plane and where possible encompassing the nuclear material within the oocyte. The following lasers were used in combination and isolation for the various staining: 488nm and 639nm. Representative images and fluorescence intensity analyses were performed on a single slice image using the Fiji software (NIH). For quantification, treatment groups were imaged on the same day using the same parameters to allow for accurate intensity comparisons. For each replicate, 5-15 individual oocytes or COCs were imaged and analysed. Thresholding (default) was performed to segment regions of interest (ROI) and analyse the mean grey value and the integrated density (sum of pixel intensity values within a ROI). For cumulus cell analysis, images containing only cumulus cells were analysed. For oocyte analysis, regions of interest (ROIs) were drawn encompassing only the oocyte for analysis. All measurements were collated in Microsoft Excel and graphs generated in Graph Pad Prism 9.5.1.

### Statistical analyses

Fertilisation rates were calculated as a proportion of the total number of oocytes that were fertilised with sperm. Hatching blastocyst and total blastocyst rates were calculated as a proportion of the total number of cleaved embryos in each treatment group. Fertilisation and blastocyst rates were represented in percentages and arcsine transformed before one-way ANOVA with Tukey’s post hoc test for statistical analysis. For fluorescence intensity, data was normalised to the mean of the *in vivo* group of each replicate before performing a one-way ANOVA with Tukey’s post hoc test for statistical analysis. No randomisation was performed on the datasets analysed in this study and confounders were not controlled. Data was analysed using Graph Pad Prism 9.5.1 and differences were considered significant when p ≤ 0.05.

## Results

### Oocyte developmental competence is compromised *in vitro*

In this study, we utilised three established models that generate oocytes of high, medium and low quality. All three models produce mature COCs at a high rate at the MII stage. However each group demonstrates variable developmental potential to form embryos, with mouse blastocyst rates of approximately 84% (*in vivo* (Sanfins, et al., 2015)), 60% (CAPA-IVM, (Zhao, et al., 2020)) and 27% (IVM, (Sanfins, et al., 2015)) in previous studies. The two *in vitro* systems examined here were CAPA-IVM (moderate grade) and standard IVM (low grade), which were compared to high grade *in vivo*-matured COCs (*in vivo*) (Figure 1).

To confirm the varying developmental potential of oocytes retrieved in our hands, we assessed embryo development through to Day 6 post-fertilisation as a direct measure of oocyte developmental competence (Figure 2A). No significant differences in 2-cell cleavage rates were observed between *in vivo*, CAPA-IVM and IVM oocyte groups at Day 2 (Figure 2B and C). Relative to *in vivo*-matured oocytes, both the CAPA-IVM and IVM groups showed lower rates of blastocyst formation by day 5 of embryo culture (Figure 2D and E), a trend which continued at day 6 (Figure 2F and H). Blastocyst hatching rates were also lower in CAPA-IVM and IVM groups relative to the *in vivo* group, by 36% and 40% respectively (Figure 2G). In all three cohorts, embryos that cleaved but did not reach the blastocyst stage were either arrested or fragmented. In summary, these data illustrates that oocytes matured *in vitro* show decreased ability to reach the blastocyst stage, and thus reduced developmental competence, when compared with oocytes matured *in vivo*.

**Figure 2.**
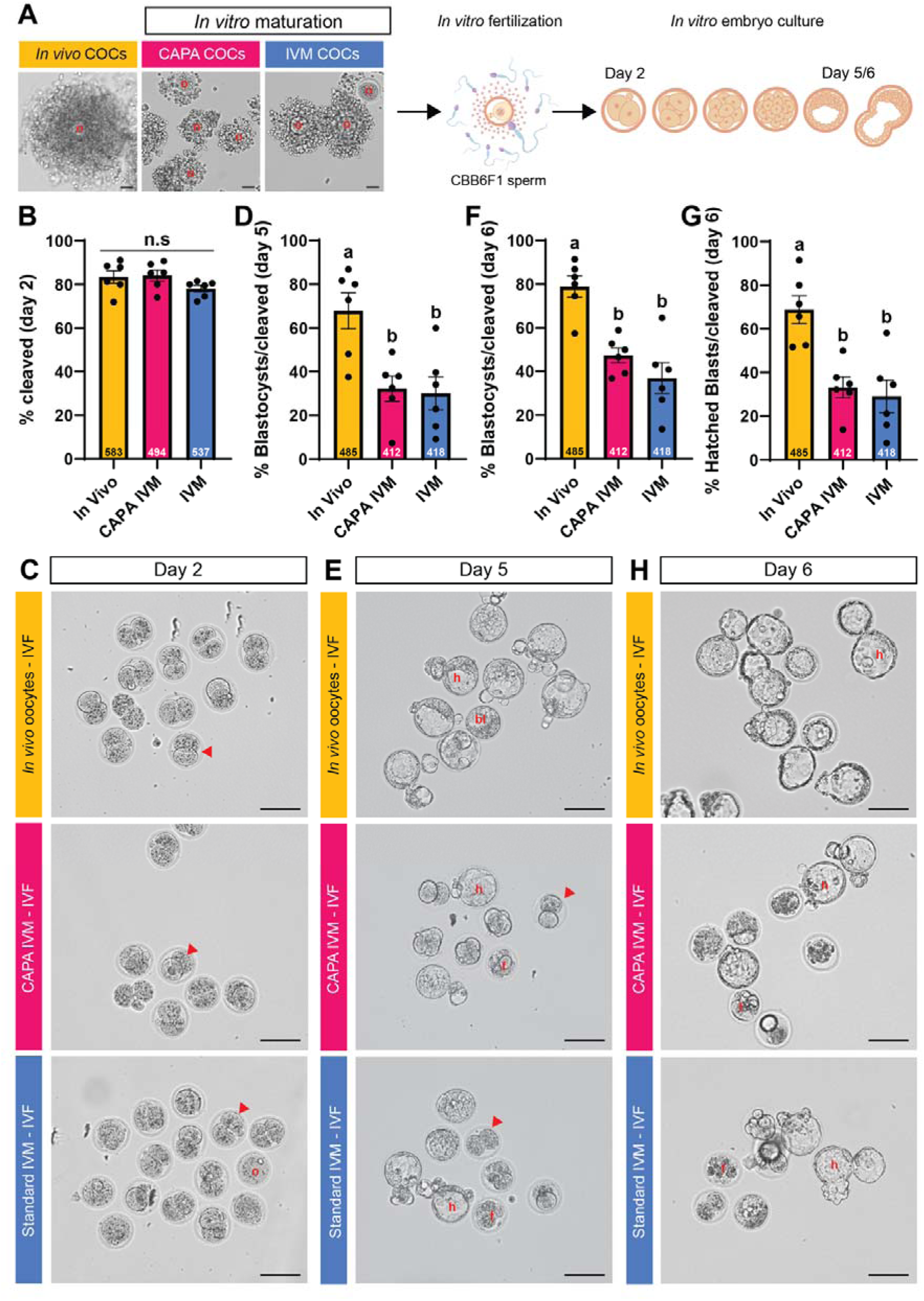
In vitro culture decreases oocyte developmental competence as measured by embryo formation and development. A) Brightfield images of cumulus-oocyte complexes (COCs) following *in vivo* maturation (yellow), CAPA IVM (pink) and standard IVM (IVM; blue). Red circles mark oocytes, and scale bar is 50 µm. Diagram of *in vitro* fertilisation (IVF) followed by morphological assessment of embryos at day 2, 5 and 6 of *in vitro* embryo culture. B) Fertilisation rate (percentage of cleaved embryos per oocyte transferred to culture), D) Day 5 blastocyst rate, F) Day 6 blastocyst rate and G) Day 6 hatching blastocyst rate (percentage of hatching blastocysts per cleaved embryo) following oocyte maturation. Corresponding images of embryos at Day 2 (C), Day 5 (E) and Day 6 (H) are shown. Red arrowheads mark cleaved embryos, red circles show oocytes that failed to cleave, hatched blastocysts (red h), blastocyst (red bl) and fragmented embryos (red f). Images taken at 10x magnification and scale bar denotes 100 µm. All embryo outcome data were arcsine transformed and one-way ANOVA followed by Tukey’s post hoc tests were performed. Numbers above the x axis denote the total number of oocytes (panel B) and total number of cleaved embryos (panel D, F, G) in each oocyte group across the six replicates. Bars with no common superscripts are significantly different (P < 0.05).

### The proteomes of both oocytes and cumulus cells are significantly altered between oocytes of high and low competence

To investigate the molecular mechanisms underpinning the variable rates of oocytes developmental competence, we used a label-free proteomic approach to comprehensively profile the proteome in oocytes and their corresponding cumulus cells. In total, we successfully identified 1,736 and 3,212 proteins in oocyte and cumulus cell samples, respectively, consistent with proteomic coverage obtained previously in other studies (Chen, et al., 2023, Richani, et al., 2024, Sun, et al., 2023, Zhang, et al., 2023) (Supplemental Datasets 1 and 2). Within the treatment groups, an average number of 8.5 (*in vivo*), 9.0 (CAPA) and 8.8 (IVM) unique peptides were identified per protein in oocytes, resulting in an average protein coverage of 27.5%, 29.2%, and 28.4%, respectively (Figure 3A). In cumulus cells, an average number of 9.6 (*In vivo*), 9.8 (CAPA) and 9.8 (IVM) unique peptides were identified per protein, encompassing an average protein coverage of 27.4%, 27.7%, and 27.9%, respectively (Figure 3B). Principal component analysis (PCA) of global proteome expression revealed diversity in oocyte replicates across principal component 1, however *in vivo*-matured oocytes separated from CAPA-IVM and standard IVM oocytes along principal component 2 (Figure 3C). Cumulus cells clustered more distinctly in PCA, with *in vivo*-matured cumulus cells segregated to the left on principal component 1 (Figure 3D). Of the proteins identified in each group (in a minimum of three replicates), most proteins were found in all treatment groups in both oocytes (86.6% overlap) and cumulus cells (92.1% overlap) (Figure 3E and F). Key oocyte proteins such as zona pellucida sperm-binding proteins 1-3 (ZP1, ZP2 and ZP3) were present in all groups, along with critical oocyte-secreted factors growth/differentiation factor 9 (GDF-9) and bone morphogenetic protein 15 (BMP-15) (Supplemental Figure 1). In cumulus cells, known marker proteins including cytochrome P450 11a1 (CYP11A1), insulin-like growth factor 1 receptor (IGF1R), pentraxin-related protein (PTX3) and prostaglandin synthase 2 (PTGS2) were present in all three groups (Supplemental Figure 1). Other cumulus cell marker proteins including steroidogenic acute regulatory protein (STAR), anti-Müllerian hormone (AMH) and FSH receptor (FSHR) were not detected in any group. Regardless, the comprehensive proteomic coverage across oocyte maturation systems allowed us to delve further into the molecular changes in both oocytes and cumulus cells.

**Figure 3.**
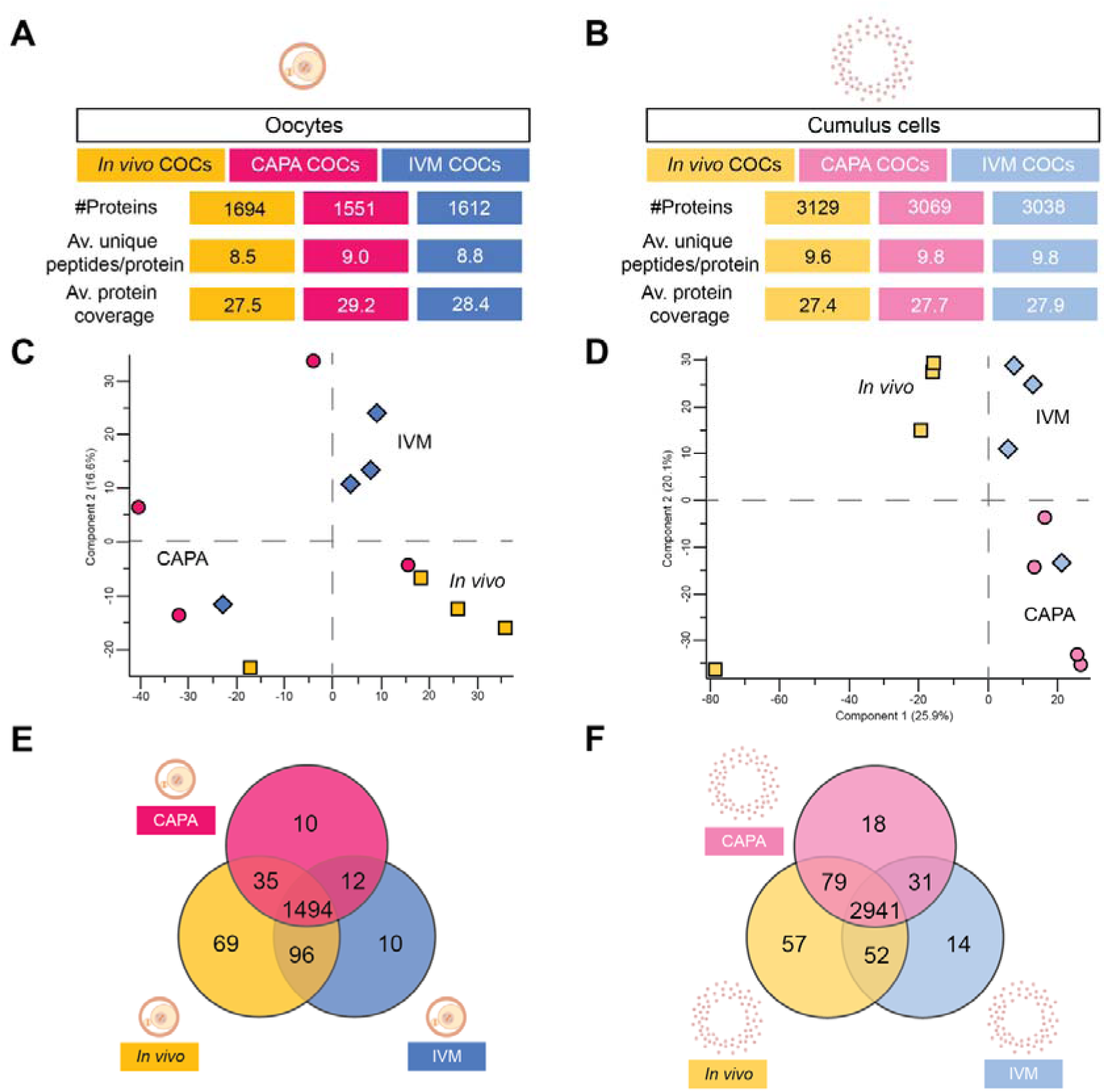
Proteomic characterisation of oocytes and cumulus cells following CAPA-IVM and standard IVM. A) The total number of proteins identified, average number of unique peptides identified/protein and the average protein coverage (%) within oocytes matured *in vivo* (yellow), following CAPA-IVM (pink) and following standard IVM (blue). B) The total number of proteins identified, average number of unique peptides identified/protein and the average protein coverage (%) within cumulus cells matured *in vivo* (light yellow), following CAPA-IVM (light pink) and following standard IVM (light blue). Principal component analysis (PCA) from each maturation method (*in vivo* (yellow squares), CAPA-IVM (pink circles) and standard IVM (blue diamonds)) for both C) oocytes and D) cumulus cells. E) Venn diagram of shared and unique proteins across the three groups of oocytes. F) Venn diagram of shared and unique proteins across the three groups of cumulus cells.

### Lower competence oocytes show expression changes in translation machinery and endocytosis proteins

Using pairwise differential expression comparisons to identify differences between oocyte cohorts, we found a small proportion of the total identified proteins (7.2% CAPA vs *in vivo*, 7.2% CAPA vs IVM and 7.5% IVM vs *in vivo*) were altered across the three models (fold-change ± 2 and p value ≤ 0.05, Figure 4A, Supplemental dataset 3). To refine our protein list, we focused on differentially expressed proteins that were dysregulated in the same direction in CAPA and IVM compared to the *in vivo* group (log2FC of ± 1, p value ≤ 0.05, Supplemental Dataset 5). This revealed a list of 34 proteins differentially expressed in both CAPA-IVM and IVM groups when compared to *in vivo* oocytes (Figure 4B). Within this list, proteins involved in eukaryotic translation (EIF2A, EIF3M, RPL24 and RPS24), RNA processing (DCP1A, HNRNPAB, HNRNPL and HNRNPU) and autophagy (CHMP2A and CHMP4B) were consistently altered. Ingenuity Pathway Analysis (IPA) of the pairwise comparisons identified several canonical pathways, with keratinisation highly enriched in CAPA-IVM oocytes (Figure 4C). In IVM oocytes, cellular stress pathways such as granzyme A signalling, DNA damage and oxidative phosphorylation were enriched when compared with *in vivo* oocytes (Figure 4D). Notably, eukaryotic translation initiation, apoptotic execution phase, meiotic synapsis and nonsense-mediated decay pathways all showed significant enrichment scores in both pairwise comparisons (Figure 4C and D).

**Figure 4.**
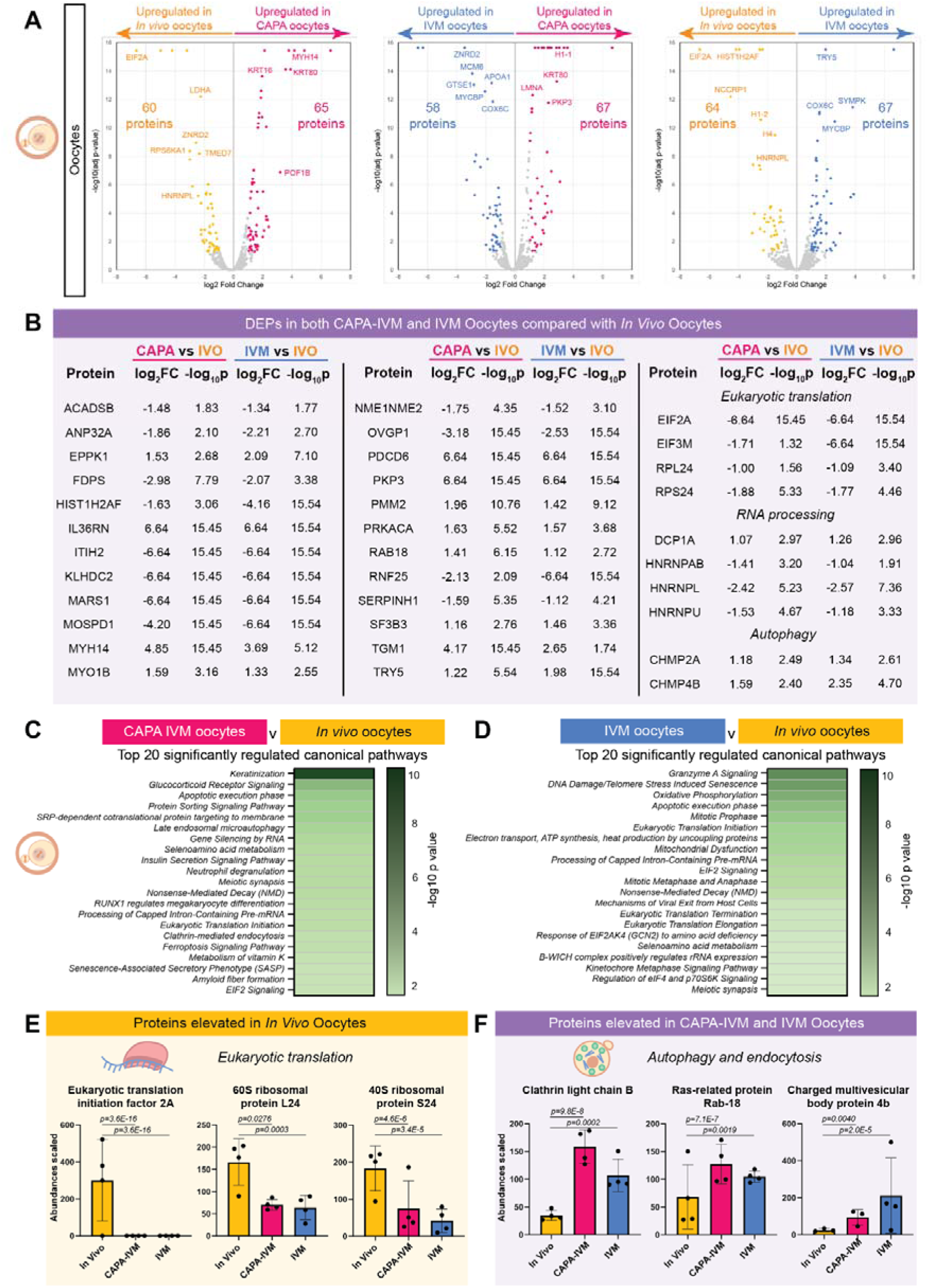
Proteomic analysis reveals disruptions in oocyte translation machinery and endocytosis proteins. A) Volcano plots showing pairwise comparisons of differential oocyte protein expression in *in vivo* (yellow), CAPA-IVM (pink) and standard IVM (blue) treatment groups. Coloured dots indicate differentially expressed proteins in each comparison and grey dots indicate proteins below the log2FC ± 1, p-value ≤ 0.05 cutoff. B) All proteins found to be consistently differentially expressed with log2 fold-change and −log10 p- values of CAPA-IVM matured MII oocytes and standard IVM matured MII oocytes compared with *in vivo* matured MII oocytes (IVO). Bar charts showing the top 20 enriched canonical pathways based on oocyte differential protein expression between C) CAPA- IVM matured MII oocytes and *in vivo* matured MII oocytes, and D) standard IVM matured MII oocytes and *in vivo* matured MII oocytes. Shades of green indicate -log10 p value generated from Ingenuity Pathway Analysis software. Scaled proteomic abundances for individual proteins related to biological pathway elevated in E) *in vivo* oocytes and F) *in vitro* matured oocytes (n = 4, abundance ratio adjusted p-values and mean ± standard deviation).

Within the proteins associated with eukaryotic translation, eukaryotic initiation factor 2A (fold change [FC] CAPA vs *in vivo* = −6.64, [FC] IVM vs *in vivo* = −6.64), 60S ribosomal protein L24 ([FC] CAPA vs *in vivo* = −1.00, [FC] IVM vs *in vivo* = −1.09) and 40S ribosomal protein S24 ([FC] CAPA vs *in vivo* = −1.88, [FC] IVM vs *in vivo* = −1.77) were all elevated in *in vivo* oocytes, with significantly lower levels in CAPA-IVM and IVM oocytes (Figure 4E). In contrast, vesicular proteins including clathrin light chain B ([FC] CAPA vs *in vivo* = 1.38, [FC] IVM vs *in vivo* = 0.94), Ras-related protein ([FC] CAPA vs *in vivo* = 1.41, [FC] IVM vs *in vivo* = 1.12) and charged multivesicular body protein 4B ([FC] CAPA vs *in vivo* = 1.59, [FC] IVM vs *in vivo* = 2.35) were highly expressed in CAPA-IVM and IVM oocytes, with lower abundances found in *in vivo* oocytes (Figure 4F). To confirm the differential expression of these target proteins across *in vivo*, CAPA-IVM and IVM oocytes, we assessed localisation of a selection of these proteins within the oocyte using immunocytochemistry.

Eukaryotic initiation factor 2A (EIF2A) showed punctate expression throughout the cytoplasm of mouse oocytes, irrespective of maturation method (Figure 5A). Expression was also observed within the polar body of all oocytes. The ribosomal subunit protein RPL24 was expressed in a diffuse pattern across the cytoplasm and polar body, with increased co-localisation with the oocyte spindle (Figure 5B). Showing similar spindle co-localisation, the endocytic protein clathrin B (CLTB) was distributed primarily to the oocyte membrane, with strong localisation to the oocyte spindle in all three treatment groups (Figure 5C). Quantification of integrated density fluorescence intensity revealed a 27% decrease in expression of EIF2A in standard IVM oocytes (but not in CAPA-IVM oocytes), when compared to *in vivo* oocytes (p=0.0025, Figure 5D). This trend was mirrored in the quantification of RPL24 intensity, where IVM oocytes, but not CAPA-IVM oocytes, demonstrated a 23% decrease in protein expression against *in vivo* oocytes (p=0.0003, Figure 5E). In contrast, CLTB was elevated by 43% in CAPA-IVM oocytes when compared to *in vivo* oocytes (p=0.0042, Figure 5F), with no significant differences observed between other treatments.

**Figure 5.**
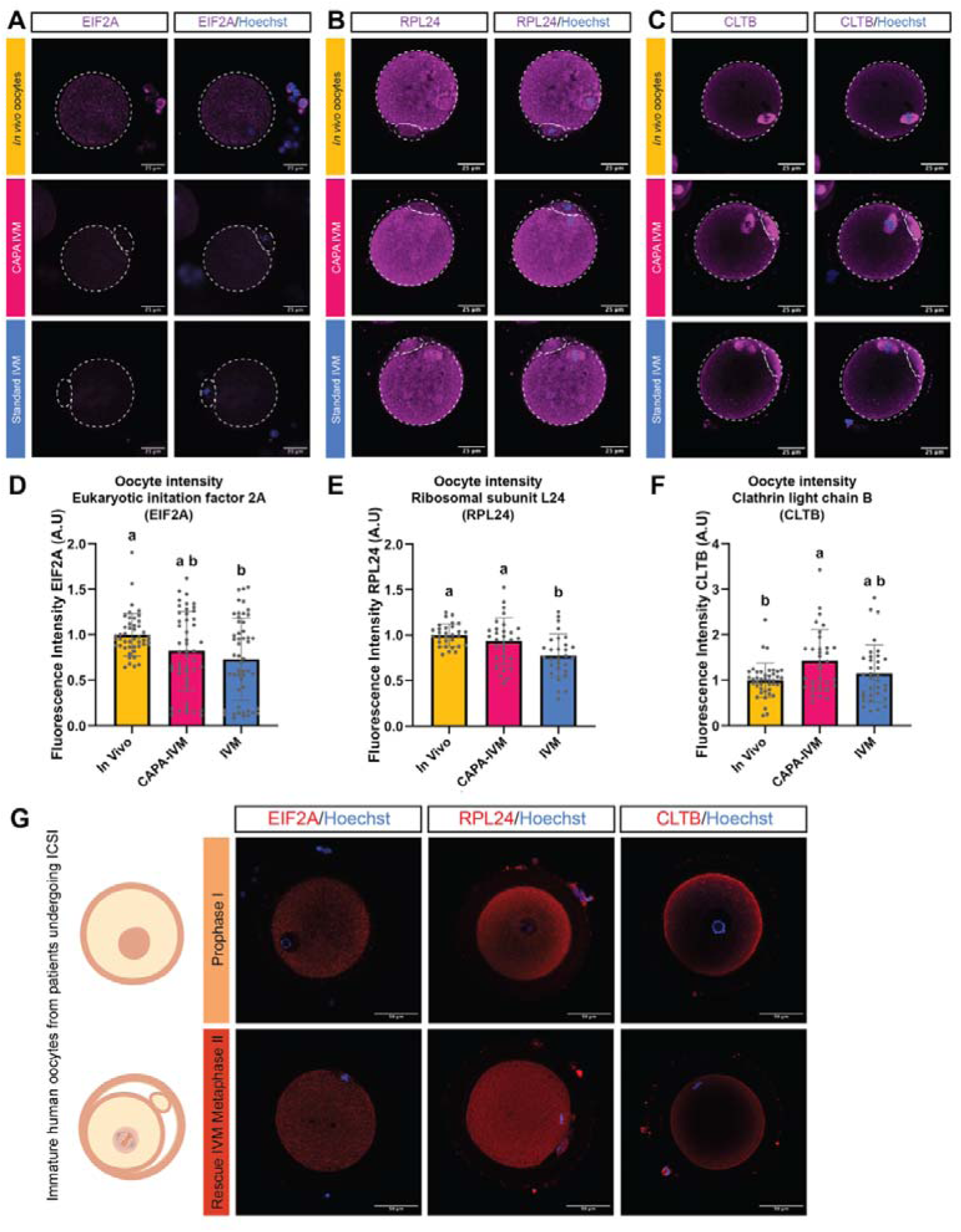
Translation machinery and endocytosis proteins are dysregulated in in vitro matured oocytes. A) Eukaryotic initiation factor 2A (EIF2A, magenta), B) ribosomal protein L24 (RPL24, magenta) and C) clathrin light chain B (CLTB, magenta) immunocytochemistry in MII oocytes following *in vivo* maturation (yellow), CAPA IVM (pink) and standard IVM (IVM; blue). Nuclei were counterstained with Hoechst (blue). Images taken at 40x magnification and scale bar denotes 25 µm. Fluorescence intensity of (D) EIF2A, (E) RPL24 and (F) CLTB was quantified in oocytes from MII COCs following *in vivo* maturation (yellow), CAPA IVM (pink) and standard IVM (IVM; blue). Integrated density values were taken from individual COCs and normalised to the *in vivo* control mean intensity of each biological replicate (dots). Data are presented as the mean of three replicates ± SEM. Statistical analyses were performed using one-way ANOVA followed by Tukey’s post hoc tests. Bars with no common superscripts are significantly different (P < 0.05). A.U, arbitrary units. G) Immunofluorescent staining for translation machinery (RPL24 and EIF2A) and endocytosis (CLTB) in human prophase I (GV) and metaphase II oocytes. Nuclei were counterstained with Hoechst (blue). Images taken at 40x magnification and scale bar denotes 50 µm.

### Proteins altered in low competence mouse oocytes are conserved in discarded human oocytes

With translation and vesicle pathway changes observed in mouse CAPA-IVM and standard IVM, we next assessed whether these differentially expressed proteins are conserved in poor-quality human GV and MII oocytes using immunocytochemistry (Figure 5G). Staining for EIF2A and RPL24 revealed conservation of these proteins in human oocytes, with EIF2A showing punctate staining across the cytoplasm and RPL24 showing diffuse staining throughout the oocyte cytoplasm (Figure 5G). No expression of either EIF2A or RPL24 was observed within the nucleolus of GV oocytes (Figure 5G). Following rescue-IVM, RPL24 and EIF2A were both expressed in MII oocytes, with staining across the cytoplasm and around the spindle, with some expression also observed in the polar body (Figure 5G). The endocytosis protein CLTB was localised to the oocyte periphery in GV oocytes, with some punctate staining within the cytoplasm (Figure 5G). This expression pattern was consistent in MII oocytes (Figure 5G). Combined, these data illustrate that key molecular pathway members EIF2A, RPL24 and CLTB identified in our mouse proteomic analysis are also present in human oocytes, highlighting the potential relevance of our developmental competence models to the human oocyte and clinical maturation strategies.

### Proteomic changes in *in vitro* cumulus cells reflects dysregulation in steroidogenic and metabolic pathways

In addition to assessing the oocyte proteome, we also profiled proteomic changes in cumulus cell proteins arising as a consequence of oocyte maturation mode. As with oocytes, pairwise differential expression comparisons revealed a small proportion of the total identified proteins were differentially expressed across the three comparisons (9.0% CAPA vs *in vivo*, 6.0% CAPA vs IVM and 7.4% IVM vs *in vivo*, log2FC of ± 1 and p value ≤ 0.05, Figure 6A, Supplemental dataset 4). We utilised the pattern-based approach to investigate the differentially expressed proteins that were altered in the same direction in CAPA-IVM and standard IVM cumulus cells (log2FC of ± 1, p value ≤ 0.05, Supplemental Dataset 6). Across the three maturation systems, 125 proteins were differentially expressed in both CAPA-IVM and IVM groups relative to the *in vivo* group (Figure 6B). This list included proteins associated with keratinisation (11 keratin proteins), which may be associated with sample contamination. The list also included a number of solute carrier proteins (SLC22A6, SLC25A30, SLC6A6, SLC7A5 and SLC7A8) (Figure 6B). To further refine our list of proteins altered *in vitro*, pathway analysis of the pairwise comparisons using IPA was performed. This revealed that the most significantly regulated canonical pathway was the response to cytosolic calcium (Figure 7A and B). Given that the calcium signalling response is well-established during oocyte maturation and fertilisation (Chen, et al., 2024), we looked for other pathways consistently altered in both comparisons. Significant canonical pathways across both comparisons included steroid biosynthesis, detoxification of reactive oxygen species, serine biosynthesis, glucocorticoid receptor signalling and cholesterol homeostasis (Figure 7A and B).

**Figure 6.**
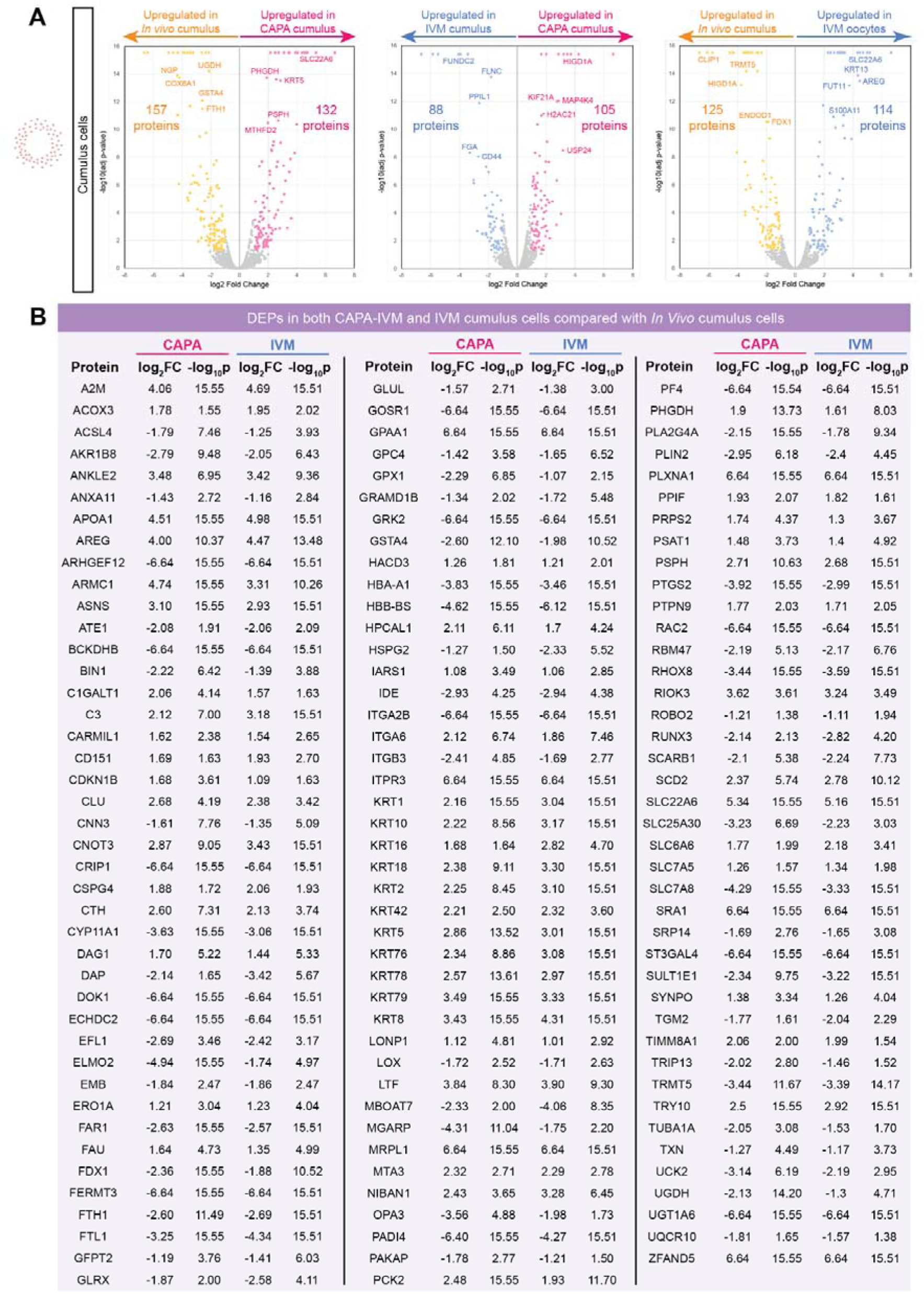
Global proteomic alterations in cumulus cells across oocyte maturation systems. A) Volcano plots showing pairwise comparisons of differential cumulus cell protein expression in *in vivo* (yellow), CAPA-IVM (pink) and IVM (blue) treatment groups. Coloured dots indicate differentially expressed proteins in each comparison and grey dots indicate proteins below the log2FC ± 1, p-value ≤ 0.05 cutoff. B) All proteins found to be consistently differentially expressed with log2 fold-change and −log10 p-values of cumulus cells from CAPA-IVM and IVM maturation groups, compared with *in vivo* matured cumulus cells (IVO).

**Figure 7.**
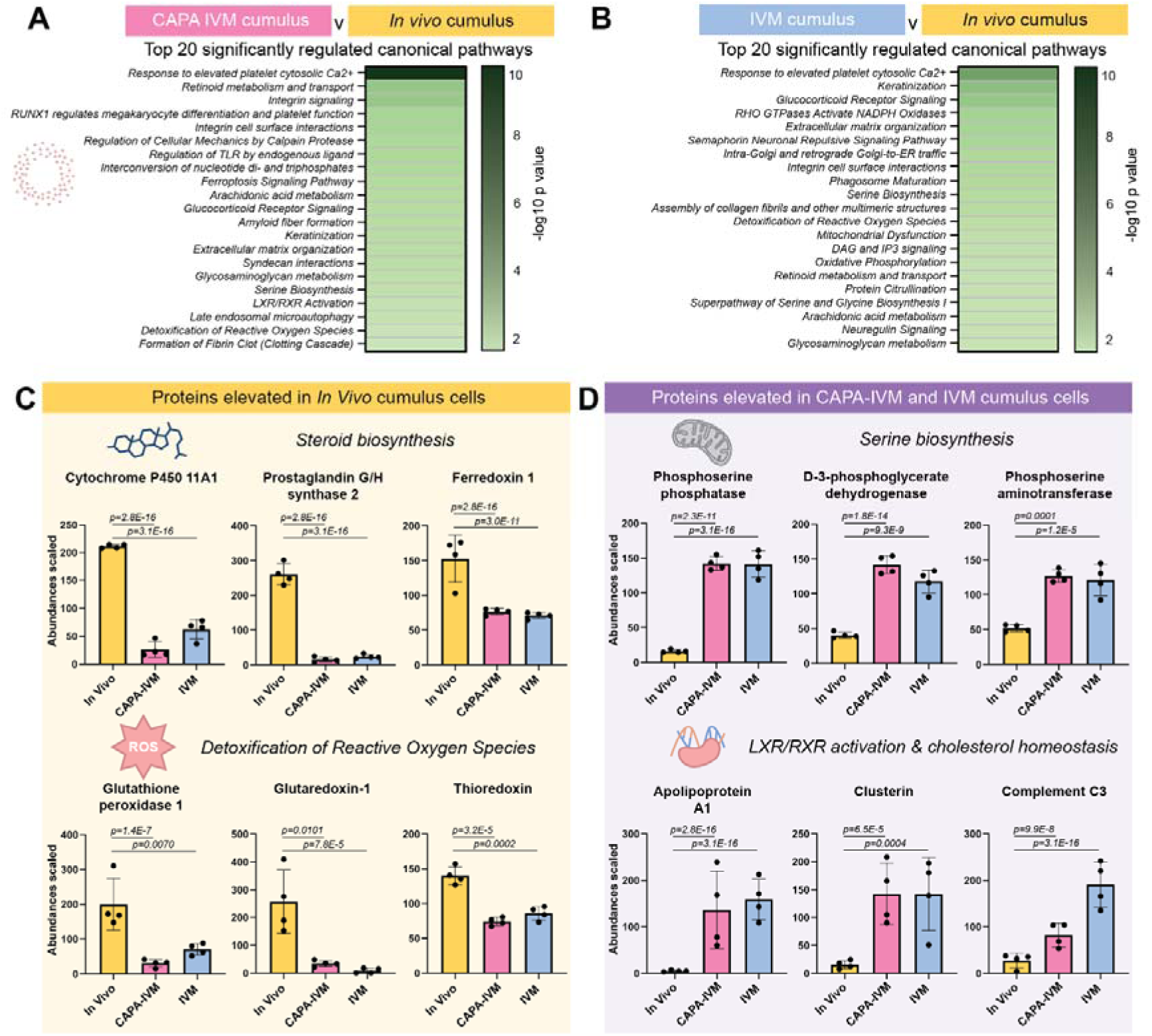
Alterations in key biological processes underpin proteomic differences in in vitro matured cumulus cells. Bar charts showing the top 20 enriched canonical pathways based on cumulus cell differential protein expression between A) CAPA-IVM matured MII COCs and *in vivo* matured MII COCs, and B) IVM matured MII COCs and *in vivo* matured MII COCs. Shades of green indicate -log10 p value generated from Ingenuity Pathway Analysis software. Scaled proteomic abundances for individual proteins related to biological pathway elevated in C) *in vivo* cumulus cells and D) *in vitro* matured cumulus cells (n = 4, abundance ratio adjusted p-values and mean ± standard deviation).

Proteins associated with these pathways showing increased expression in *in vivo* cumulus cells included cytochrome P450 11A1 (fold change [FC] CAPA vs *in vivo* = −3.63, [FC] IVM vs *in vivo* = −3.06), prostaglandin G/H synthase 2 ([FC] CAPA vs *in vivo* = −3.92, [FC] IVM vs *in vivo* = −2.99) and ferredoxin 1 ([FC] CAPA vs *in vivo* = −2.36, [FC] IVM vs *in vivo* = −1.88); all key proteins involved in the critical function of steroid biosynthesis in cumulus cells (Figure 7C). Additionally, proteins associated with cellular anti-oxidant response, glutathione peroxidase 1 ([FC] CAPA vs *in vivo* = −2.29, [FC] IVM vs *in vivo* = −1.07), glutaredoxin 1 ([FC] CAPA vs *in vivo* = −1.87, [FC] IVM vs *in vivo* = −2.58), and thioredoxin ([FC] CAPA vs *in vivo* = −1.87, [FC] IVM vs *in vivo* = −2.58), showed significantly higher expression in *in vivo* cumulus cells compared to CAPA-IVM and IVM cumulus cells (Figure 7C). Conversely, proteins associated with serine and cholesterol biosynthesis were aberrantly elevated in CAPA-IVM and IVM cumulus cells when compared to *in vivo* cumulus cells (Figure 7D). These proteins included phosphoserine phosphatase ([FC] CAPA vs *in vivo* = 2.71, [FC] IVM vs *in vivo* = 2.68), D-3 phosphoglycerate dehydrogenase ([FC] CAPA vs *in vivo* = 1.90, [FC] IVM vs *in vivo* = 1.61), phosphoserine aminotransferase ([FC] CAPA vs *in vivo* = 1.48, [FC] IVM vs *in vivo* = 1.40), apolipoprotein A1 ([FC] CAPA vs *in vivo* = 4.51, [FC] IVM vs *in vivo* = 4.98), clusterin ([FC] CAPA vs *in vivo* = 2.68, [FC] IVM vs *in vivo* = 2.38) and complement C3 ([FC] CAPA vs *in vivo* = 2.12, [FC] IVM vs *in vivo* = 3.18).

Given that both steroid production and metabolic homeostasis are essential components of cumulus cell function, particularly during oocyte maturation (Chen, et al., 2023, Richani, et al., 2021), we then focused on characterising target proteins within the cumulus cells in each maturation strategy using immunocytochemistry (Figure 8A and C). Ferredoxin 1 (FDX1) was intensely expressed in the cytoplasm of expanded *in vivo* cumulus cells, colocalising with the actin cytoskeleton stain phalloidin (Figure 8A). In CAPA and IVM cumulus cells, which demonstrate a decreased ability to expand in culture, FDX1 expression was variable, where some but not all sections of the cumulus cell clusters showed high expression (Figure 8A). Quantification of all three groups revealed a significant decline in fluorescence intensity *in vitro*, with a 58% decrease in CAPA-IVM cumulus cells and a 65% decrease in standard IVM cumulus cells, when compared to *in vivo* cumulus cells (Figure 8B). This analysis confirmed the decline in FDX1 protein levels observed by mass spectrometry, validating altered steroid biosynthesis in cumulus cells matured *in vitro* (Figure 7C). We then examined D-3 phosphoglycerate dehydrogenase (PHGDH) using the same approach and found low expression of PHGDH in *in vivo* cumulus cells (Figure 8C). Levels of PHGDH were elevated in CAPA-IVM cumulus cells throughout the nucleus and cytoplasm, with variable expression observed in standard IVM cumulus cells (Figure 8D). CAPA-IVM cumulus cell intensity levels of PHGDH were increased 160% and 99% compared to *in vivo* and standard IVM cumulus cells respectively, which may indicate that CAPA-IVM cumulus cells have a higher requirement for components of the serine biosynthesis pathway. Combined, the data confirms that alterations observed in the proteomic analysis are reflective of oocyte quality, both in mechanisms in the oocyte and their accompanying cumulus cells.

**Figure 8.**
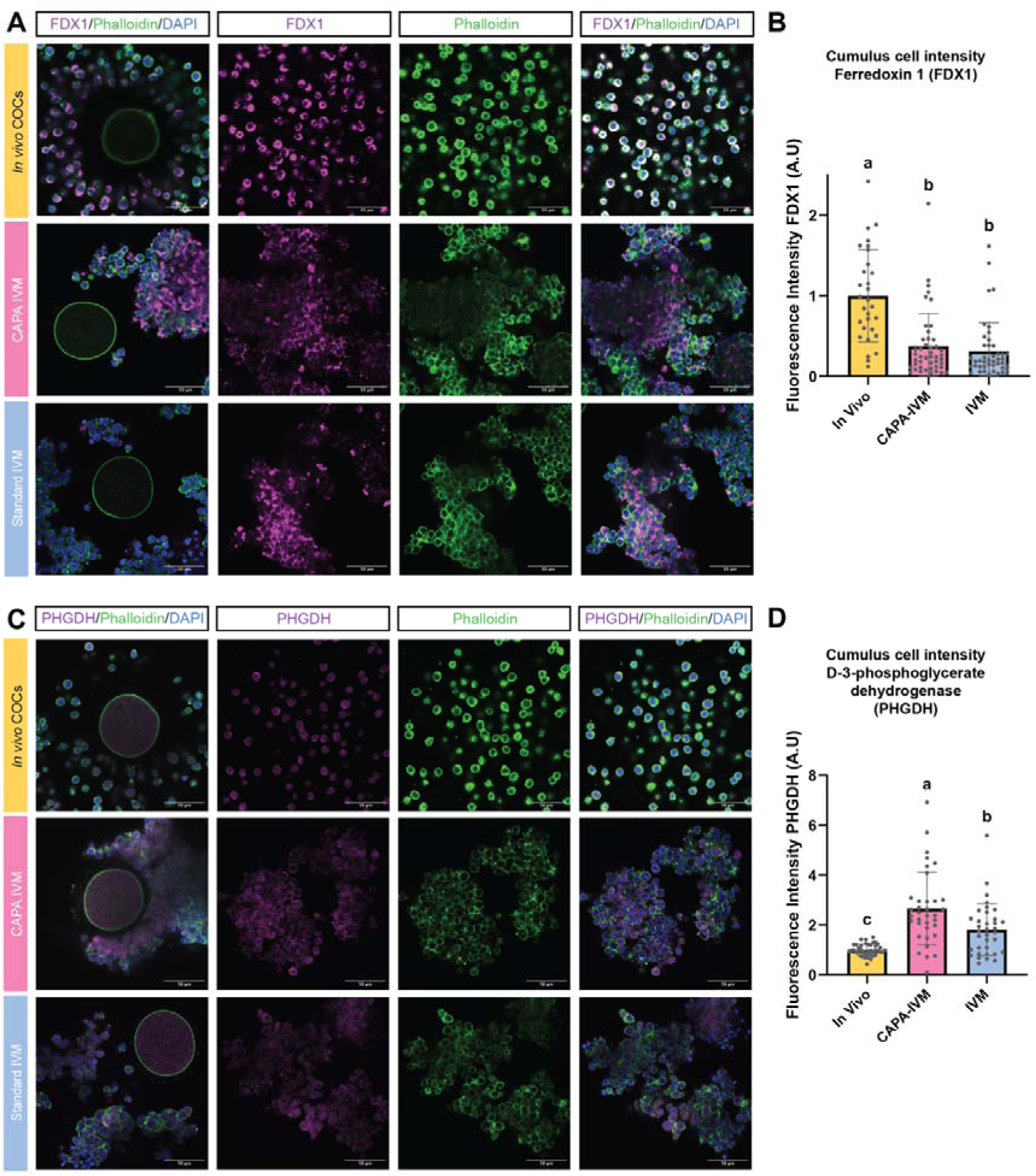
Mitochondrial protein FDX1 and serine biosynthesis enzyme PHGDH are dysregulated in in vitro matured cumulus cells. A) Ferredoxin 1 (FDX1, magenta) and C) D-3-phosphoglycerate dehydrogenase (PHGDH, magenta) immunocytochemistry colocalised with phalloidin (green) in MII COCs following *in vivo* maturation (yellow), CAPA IVM (pink) and standard IVM (IVM; blue). Nuclei were counterstained with Hoechst (blue). First column shows intact cumulus oocyte complexes, with the subsequent columns showing representative images of cumulus cells. Images taken at 40x magnification and scale bar denotes 50 µm. Fluorescence intensity of (B) FDX1 and (D) PHGDH was quantified in cumulus cells from MII COCs following *in vivo* maturation (yellow), CAPA IVM (pink) and standard IVM (IVM; blue). Integrated density values were taken from individual COCs and normalised to the *in vivo* control mean intensity of each biological replicate (dots). Data are presented as the mean of three replicates ± SEM. Statistical analyses were performed using one-way ANOVA followed by Tukey’s post hoc tests. Bars with no common superscripts are significantly different (P < 0.05). A.U, arbitrary units.

## Discussion

Oocyte quality is a critical foundation for reproductive success. Despite learnings from ART and decades of research in multiple species, we still lack detailed knowledge of the complex biological processes that underpin oocyte quality, and thus future potential to support healthy embryo development. In this study, we assessed the molecular pathways underpinning developmental competence across mouse MII cumulus-oocyte complexes, that had undergone clinically relevant maturation protocols, and checked conservation of target proteins in human oocytes. Proteomic profiling revealed global alterations in both oocytes and cumulus cells between high and low competence mouse COCs, revealing a plethora of novel biological processes that may contribute to oocyte developmental competence. Orthogonal approaches confirmed aberrant changes to translation machinery and endocytosis proteins in oocytes following CAPA-IVM and IVM, with more profound changes in cumulus cell pathways including steroid, serine and cholesterol biosynthesis proteins. Combined, these data shed new light on the mechanisms contributing to the varied developmental competence seen in pre-IVM and IVM systems, which have important clinical and biological consequences for our understanding and assessment of oocyte quality.

The focus of clinical ART improvement has predominately centred on optimising ovarian stimulation using gonadotrophins (Schouten, et al., 2025). For the sub-set of ART patients that are particularly sensitive to gonadotrophins, for example women with polycystic ovarian syndrome (PCOS), IVM represents an alternative, safe and low-intervention treatment option (Gilchrist and Smitz, 2023). Recently, the shift towards revisiting IVM protocols and creating new reproductive technologies have been aided by the surge of basic knowledge in oocyte and ovarian biology. CAPA-IVM, a biphasic form of *in vitro* maturation, relies on CNP regulating meiotic arrest through signalling from the cumulus cells (Sanchez, et al., 2019). CAPA-IVM emerged from a number of differing pre-IVM systems, designed to improve oocyte quality through maintained oocyte-cumulus cell communication and extended time for the accumulation of important molecular machinery (Gilchrist, et al., 2024). In this study, we modelled our CAPA-IVM and standard IVM treatment timings on established clinical IVM protocols (De Vos, et al., 2025, Vuong, et al., 2020a), where mice received minimal stimulation in the form of a short 23 hr PMSG priming, generating intermediate and moderate-competence oocytes respectively (Nogueira, et al., 2003, Zhao, et al., 2020). While we did not observe a statistical difference in embryo outcomes between CAPA-IVM and standard IVM, as others have shown in mice and humans (Sánchez, et al., 2017, Vuong, et al., 2020b, Zhao, et al., 2020), our proteomics data shows comprehensive changes in identified proteins and differences in expression levels between CAPA-IVM and standard IVM cumulus-oocyte complexes. This is in line with previous studies revealing molecular alterations in CAPA-IVM mouse oocytes, improving their quality (Akin, et al., 2021, Zhao, et al., 2020). Of greater clinical interest are the substantial proteomic differences observed between high competence oocytes matured *in vivo*, compared to lower competence oocytes matured *in vitro* by either CAPA-IVM or standard IVM. While CAPA-IVM and standard IVM are well-suited to certain patient populations, such as those with PCOS, the cumulative live birth rate following CAPA-IVM remains lower than conventional IVF (Nguyen, et al., 2022). With the increased uptake and emerging interest in pre-IVM systems, particularly CAPA-IVM, worldwide, there exists an opportunity to further capitalise on reproductive biology research to precisely enhance *in vitro* protocols, in addition to meiotic arrest.

The proteomic dataset generated in this manuscript provides an unbiased approach to assessing pathways that might dictate oocyte developmental competence. It supplements other mouse studies analysing the metabolome (Chen, et al., 2023), transcriptome (Gao, et al., 2017, Ma, et al., 2013, Vigone, et al., 2013), epigenome (Wasserzug Pash, et al., 2022), proteome (Richani, et al., 2024), secretome (Fiorentino, et al., 2024), active translation (Dvoran, et al., 2024) and the chromatin landscape (Zhu, et al., 2025) of mouse oocytes and/or cumulus cells. Given that the mature oocyte is largely considered transcriptionally quiescent, proteomic analysis provides the link between the stored mRNA of the oocyte and the dynamic actions of molecular machinery in actioning necessary cellular processes. Nevertheless, consolidating our proteomics dataset with these previously published studies of *in vivo* and IVM mouse oocytes will create a more comprehensive network of oocyte developmental competence at the MII stage. This will be necessary to untangle the expression of keratin proteins in our *in vitro* oocyte groups. Keratins are essential during early pre-implantation embryo development (Lim, et al., 2020) and may be acquired by the oocyte as a component of developmental competence, but could also be explained by the increased handling and culture periods of *in vitro* matured oocytes.

Paired analyses of the oocyte and its corresponding cumulus cells are also critical to understanding developmental competence. Not only do the oocytes produce and store biochemical components themselves, but increasing evidence suggests that cumulus cells will deplete their stores and donate metabolites, small molecules and possibly even RNA to the oocyte (Doherty, et al., 2025, Macaulay, et al., 2014, Richani, et al., 2019). Analysing both the oocyte and cumulus cells separately in future studies will allow for in-depth interrogation of the substrates donated to the oocyte across the course of *in vitro* maturation. Potential opportunities also exist to test conservation of our novel pathways in human oocytes, with the caveat that most research material is performed on oocytes that are deemed developmentally incompetent and thus discarded. Given the conservation already observed across species for other oocyte-specific molecular processes (Pek, 2024), other aspects of developmental competence are likely to be evolutionarily essential.

The field of oocyte developmental competence is rapidly expanding (for detailed reviews, see (Conti and Franciosi, 2018, Richani and Gilchrist, 2021, Tan and Dunning, 2022)). In addition, it is now appreciated that both nuclear and cytoplasmic maturation are inextricably linked and must be synchronised to ensure the best gamete potential (Eppig, 1996). Within the concept of developmental competence, the oocyte holds many unusual features, for example, the presence of a midbody cap (Jung, et al., 2023), exit from electron transport chain dormancy (Rodríguez-Nuevo, et al., 2022), degradative waste processes (Zaffagnini, et al., 2024) and cytoplasmic lattices for protein storage (Jentoft, et al., 2023, Williams, et al., 2025), all key components of a mature oocyte competent for fertilisation. Within our proteomic lists, oocyte proteins involved in zona pellucida formation (ZP1-ZP3), bi-directional communication (GDF9 and BMP15), cytoplasmic scaffolding (PADI6) and vesicular encapsulation (RUFY1) are all consistently expressed across *in vivo*, CAPA-IVM and IVM maturation strategies. This illustrates that oocytes matured by all three methods would be deemed as high-quality by all previous demonstrations of high developmental competence, thus highlighting the need for new indicators of gamete quality within a mature oocyte.

To achieve developmental competence, oocytes must remodel their proteome to not only acquire the maternal transcripts and proteins to last through the maternal-zygotic transition but must also degrade proteins that are no longer required. Specific maternal transcripts are targeted and degraded across the GV-to-MII transition, including ribosomal protein subunits (Su, et al., 2007). Our proteomics data showing alterations in eukaryotic translation proteins could be explained by either of two lines of reasoning: 1) that aberrant degradation of eukaryotic translation factor and ribosomal protein subunits occur in CAPA-IVM and IVM oocytes resulting in lower protein abundances, or 2) that these proteins are already lower in abundance due to the shortened stimulation period of PMSG, and thus show lower abundance following CAPA-IVM and IVM. Regardless, promoting the appropriate removal of transcripts and machinery is a critical requirement within pre-IVM, and will ensure integrity of the oocyte translational programme.

In addition to maternal transcript degradation, redundant proteins must also be tagged and processed for waste removal or amino acid recycling. Oocytes have developed unusual vesicular strategies, termed endo-lysosomal vesicular assemblies (ELVAs) to sequester protein waste within the oocyte, observed in both mice and humans (Zaffagnini, et al., 2024, Zaffagnini, et al., 2025). In this study, proteins involved with autophagy and endocytosis (CLTB, RAB18 and CHMP4A) were differentially upregulated in CAPA-IVM and IVM oocytes, when compared to *in vivo* oocytes, suggesting an accumulation of vesicles and their contents *in vitro* (Figure 4E). While the identification of ELVAs is recent, one study assessing macroautophagy found an accumulation of LAMP1-positive vesicles in the GV oocytes of reproductively aged mice, known to have poor oocyte quality (Peters, et al., 2021). This mechanism may be conserved in our systems of low developmental competence, suggesting a link between the molecular changes occurring with ageing and *in vitro* maturation. Importantly, many proteins involved in vesicular processes, including clathrin and dynamin, also co-localise to the oocyte spindle, suggesting a two-pronged approach of this cellular machinery in promoting oocyte developmental competence (Camia, et al., 2025, Mihalas, et al., 2020). In this study, we observed no clear change in the localisation of clathrin light chain B to the oocyte membrane and spindle within *in vivo*, CAPA-IVM or IVM oocytes (Figure 5C). However, closer analysis of clathrin localisation and intensity on the oocyte spindle and across spindle assembly using live imaging may reveal changes across the GV-to-MII transition in CAPA-IVM and IVM oocytes.

Translating results from mouse models into the human clinical setting presents a challenge, given that good quality human oocytes are exceptionally rare research material. However, assessing the conservation of molecular pathways is essential, as often molecular strategies adopted by oocytes vary between species (Pek, 2024). Recent work has highlighted proteostasis and lysosome activity are reduced across the GV-MII translation in human oocytes, in contrast to findings in mice (Zaffagnini, et al., 2024, Zaffagnini, et al., 2025). Our demonstration that key eukaryotic translation and endocytosis proteins EIF2A, RPL24 and CLTB are expressed within human oocytes presents an exciting finding. While the assessment of three key proteins selected in this study is descriptive, opportunities exist to compare discarded human oocytes from traditional IVF cycles with oocytes from CAPA-IVM treated patients. This would confirm if the trends observed in our mouse models hold true for human oocytes. Using a similar approach to this study combining label-free proteomics and immunofluorescence staining in human oocytes may reveal conserved factors critical for oocyte quality, resulting in new oocyte quality assessments or targeted additives that can improve pregnancy outcomes for patients with poor quality oocytes.

Our proteomics data confirmed that comprehensive remodelling of the cumulus cell proteome occurs in the absence of the follicular microenvironment. More proteins were differentially expressed in comparisons between *in vivo* cumulus cells and the *in vitro* matured cumulus cells, reflective of poor cumulus cell expansion observed *in vitro* and that the cumulus cell proteome is highly susceptible to proteomic changes once removed from the follicular microenvironment. It is important to note that our dataset is limited by an inability to control exposure to the LH cascade in our *in vivo* group, necessary for producing ovulated MII oocytes. Regardless, established pathways including steroid production and glutathione detoxification were altered *in vitro*; an additional confirmation of their established functions in cumulus cells for producing high quality oocytes and subsequent embryos (Maedomari, et al., 2007, Trapphoff, et al., 2016). The identification of elevated ferredoxin-1 in *in vivo* cumulus cells links changes in steroidogenic and metabolic homeostasis, as FDX1 is an iron-sulphur protein that indirectly transfer electrons from NADPH to mitochondrial cytochrome p450 enzymes (Xing, et al., 2023). In contrast, the increase in proteins involved in serine biosynthesis in both CAPA-IVM and IVM cumulus cells, suggest an imbalance in amino acid production and a potential compensatory effect. Increased serine biosynthesis has been observed in the cumulus cells of women with diminished ovarian reserve, associating serine synthesis proteins with ovarian dysfunction (Lu, et al., 2023). Combined, our data builds on the expanding evidence that metabolic homeostasis and mitochondrial function in cumulus cells is essential to enhanced oocyte developmental competence (Akin, et al., 2021, Chen, et al., 2023, Herta, et al., 2023, Richani, et al., 2024, Tan, et al., 2022).

In summary, we have compared three models of oocyte maturational competence, showing differential proteomic alterations in oocytes and cumulus cells matured *in vitro*, relative to those matured *in vivo*. Compromised translation and vesicular processes in *in vitro* oocytes, along with functional changes in metabolic and amino acid production within cumulus cells, are promising new contributors to oocyte developmental competence. As the global ART pregnancy rate plateaus (Chambers, et al., 2021), new insights into oocyte developmental competence holds the potential to improve clinical oocyte diagnostics, expand on current clinical maturation systems and provide novel therapeutic options to infertile patients. With the emerging pre-IVM and IVM treatment options, a strengthened understanding of oocyte developmental competence will improve the developmental capacity of *in vitro* matured oocytes. This research builds towards the overall clinical goal of ART, to achieve high-quality oocytes and eventual offspring for patients, regardless of the mode of oocyte maturation.

## Supplemental data

This article contains supplemental data (1 figure, 2 tables and 6 datasets).

## Data availability

The mass spectrometry proteomics data have been deposited to the ProteomeXchange Consortium via the PRIDE (Perez-Riverol, et al., 2025) partner repository with the dataset identifier PXD073269 https://doi.org/10.6019/PXD073269.

## Acknowledgments

The authors gratefully acknowledge the team of clinicians and laboratory staff in the Fertility & Research Centre at the Royal Hospital for Women Sydney for collecting the human oocytes using in this study. Mass spectrometric results were obtained at the Bioanalytical Mass Spectrometry facility within the Mark Wainwright Analytical Centre of UNSW. Imaging for this study was performed using instruments housed in and maintained by the Biomedical Imaging Facility (KGLMF) at UNSW. The authors also acknowledge David Skerrett-Byrne for assistance and discussions with the data analysis performed in this manuscript.

## Authors’ roles

E.R.F., D.R and R.B.G conceived the study. E.R.F wrote the manuscript with help from R.B.G. E.R.F, A.P, B.P.M, I.E.S, L.E.W and R.B.G provided expertise and feedback on the study design. All experiments were performed by E.R.F, D.R, A.P, X.L, E.G, B.G, A.B, A.V, B.P.M, I.E.S and K.K. Mass spectrometry experiments were performed by A.P and analysis was performed by E.R.F. All the authors critically revised the manuscript. R.B.G provided resources for this work.

## References

Akin N, von Mengden L, Herta AC, Billooye K, van Leersum J, Cava-Cami B, Saucedo-Cuevas L, Klamt F, Smitz J, Anckaert E. Glucose metabolism characterization during mouse *in vitro* maturation identifies alterations in cumulus cellsdagger. Biol Reprod 2021;104: 902–913.

Camia B, Longo M, Bergonzi A, Dezza I, Biggiogera M, Redi CA, Casasco A, Monti M. The localization and function of the moonlighting protein Clathrin during oocyte maturation. Developmental biology 2025;517: 1–12.

Chambers GM, Dyer S, Zegers-Hochschild F, de Mouzon J, Ishihara O, Banker M, Mansour R, Kupka MS, Adamson GD. International Committee for Monitoring Assisted Reproductive Technologies world report: assisted reproductive technology, 2014†. Human reproduction (Oxford, England) 2021;36: 2921-2934.

Chen C, Huang Z, Dong S, Ding M, Li J, Wang M, Zeng X, Zhang X, Sun X. Calcium signaling in oocyte quality and functionality and its application. Frontiers in endocrinology 2024;15: 1411000.

Chen M, Yang W, Guo Y, Hou X, Zhu S, Sun H, Guo X, Chen M, Wang Q. Multi-omics reveal the metabolic patterns in mouse cumulus cells during oocyte maturation. Journal of ovarian research 2023;16: 156.

Conti M, Franciosi F. Acquisition of oocyte competence to develop as an embryo: integrated nuclear and cytoplasmic events. Human Reproduction Update 2018;24: 245–266.

De Vos M, Mostinckx L, Drakopoulos P, Anckaert E, Smitz J, Mackens S, Blockeel C, Segers I. Randomized controlled trial to evaluate the impact of follicle priming on IVM outcomes in women with polycystic ovaries: CFA versus FSH-B. Human Reproduction 2025;40: 1127–1137.

Doherty CA, Tijjani A, Munger SC, Laird DJ. Mammalian oocytes receive maternal-effect RNAs from granulosa cells. bioRxiv 2025: 2025.2002.2010.637575.

Dvoran M, Iyyappan R, Masek T, Pospisek M, Kubelka M, Susor A. Assessment of active translation in cumulus-enclosed and denuded oocytes during standard *in vitro* maturation and early embryo development. Human Reproduction 2024.

Eppig JJ. Coordination of nuclear and cytoplasmic oocyte maturation in eutherian mammals. Reproduction, fertility, and development 1996;8: 485–489.

Fatehi AN, Zeinstra EC, Kooij RV, Colenbrander B, Bevers MM. Effect of cumulus cell removal of *in vitro* matured bovine oocytes prior to *in vitro* fertilization on subsequent cleavage rate. Theriogenology 2002;57: 1347–1355.

Fiorentino G, Merico V, Zanoni M, Comincini S, Sproviero D, Garofalo M, Gagliardi S, Cereda C, Lin C-J, Innocenti F et al. Extracellular vesicles secreted by cumulus cells contain microRNAs that are potential regulatory factors of mouse oocyte developmental competence. Mol Hum Reprod 2024;30.

Gao L, Jia G, Li A, Ma H, Huang Z, Zhu S, Hou Y, Fu X. RNA-Seq transcriptome profiling of mouse oocytes after *in vitro* maturation and/or vitrification. Scientific reports 2017;7: 13245.

Geshi M, Takenouchi N, Yamauchi N, Nagai T. Effects of sodium pyruvate in nonserum maturation medium on maturation, fertilization, and subsequent development of bovine oocytes with or without cumulus cells. Biol Reprod 2000;63: 1730–1734.

Gilchrist RB, Ho TM, De Vos M, Sanchez F, Romero S, Ledger WL, Anckaert E, Vuong LN, Smitz J. A fresh start for IVM: capacitating the oocyte for development using pre-IVM. Hum Reprod Update 2024;30: 3–25.

Gilchrist RB, Smitz J. Oocyte *in vitro* maturation: physiological basis and application to clinical practice. Fertility and sterility 2023;119: 524–539.

Hashimoto S, Saeki K, Nagao Y, Minami N, Yamada M, Utsumi K. Effects of cumulus cell density during *in vitro* maturation of the developmental competence of bovine oocytes. Theriogenology 1998;49: 1451–1463.

Herta AC, von Mengden L, Akin N, Billooye K, Coucke W, Cava-Cami B, Klamt F, Smitz J, Anckaert E. Glucose and redox metabolism in meiotically blocked *in vitro* grown mouse antral follicles. Journal of assisted reproduction and genetics 2023;40: 2851–2863.

Jentoft IMA, Bäuerlein FJB, Welp LM, Cooper BH, Petrovic A, So C, Penir SM, Politi AZ, Horokhovskyi Y, Takala I et al. Mammalian oocytes store proteins for the early embryo on cytoplasmic lattices. Cell 2023;186: 5308–5327.e5325.

Jung GI, Londono-Vasquez D, Park S, Skop AR, Balboula AZ, Schindler K. An oocyte meiotic midbody cap is required for developmental competence in mice. Nature communications 2023;14: 7419.

Krämer A, Green J, Pollard J, Jr., Tugendreich S. Causal analysis approaches in Ingenuity Pathway Analysis. Bioinformatics 2014;30: 523–530.

Lemseffer Y, Terret ME, Campillo C, Labrune E. Methods for Assessing Oocyte Quality: A Review of Literature. Biomedicines 2022;10.

Li L, Zheng P, Dean J. Maternal control of early mouse development. Development 2010;137: 859–870.

Lim HYG, Alvarez YD, Gasnier M, Wang Y, Tetlak P, Bissiere S, Wang H, Biro M, Plachta N. Keratins are asymmetrically inherited fate determinants in the mammalian embryo. Nature 2020;585: 404–409.

Lu X, Lv X, Dong X, Li Y, Turathum B, Liu S, Wang X, Shi H, Liu Y. Increased serine synthesis in cumulus cells of young infertile women with diminished ovarian reserve. Human Reproduction 2023;38: 1723–1732.

Ma JY, Li M, Luo YB, Song S, Tian D, Yang J, Zhang B, Hou Y, Schatten H, Liu Z et al. Maternal factors required for oocyte developmental competence in mice: transcriptome analysis of non- surrounded nucleolus (NSN) and surrounded nucleolus (SN) oocytes. Cell Cycle 2013;12: 1928–1938.

Macaulay AD, Gilbert I, Caballero J, Barreto R, Fournier E, Tossou P, Sirard MA, Clarke HJ, Khandjian É W, Richard FJ et al. The gametic synapse: RNA transfer to the bovine oocyte. Biol Reprod 2014;91: 90.

Maedomari N, Kikuchi K, Ozawa M, Noguchi J, Kaneko H, Ohnuma K, Nakai M, Shino M, Nagai T, Kashiwazaki N. Cytoplasmic glutathione regulated by cumulus cells during porcine oocyte maturation affects fertilization and embryonic development *in vitro*. Theriogenology 2007;67: 983–993.

Mihalas BP, Pieper GH, Aboelenain M, Munro L, Srsen V, Currie CE, Kelly DA, Hartshorne GM, Telfer EE, McAinsh AD et al. Age-dependent loss of cohesion protection in human oocytes. Current biology : CB 2024;34: 117–131.e115.

Mihalas BP, Redgrove KA, Bernstein IR, Robertson MJ, McCluskey A, Nixon B, Holt JE, McLaughlin EA, Sutherland JM. Dynamin 2-dependent endocytosis is essential for mouse oocyte development and fertility. FASEB J 2020;34: 5162–5177.

Nguyen DL, Nguyen NA, Pham TD, Nguyen MHN, Vuong LN. Development of children born after *in vitro* maturation with a prematuration step versus natural conception: a prospective cohort study. Journal of assisted reproduction and genetics 2022;39: 1959–1965.

Nogueira D, Cortvrindt R, De Matos DG, Vanhoutte L, Smitz J. Effect of phosphodiesterase type 3 inhibitor on developmental competence of immature mouse oocytes *in vitro*. Biol Reprod 2003;69: 2045–2052.

Pek JW. The idiosyncrasies of oocytes. Trends Cell Biol 2024.

Perez-Riverol Y, Bandla C, Kundu DJ, Kamatchinathan S, Bai J, Hewapathirana S, John NS, Prakash A, Walzer M, Wang S et al. The PRIDE database at 20 years: 2025 update. Nucleic acids research 2025;53: D543–d553.

Peters AE, Caban SJ, McLaughlin EA, Roman SD, Bromfield EG, Nixon B, Sutherland JM. The Impact of Aging on Macroautophagy in the Pre-ovulatory Mouse Oocyte. Frontiers in cell and developmental biology 2021;9: 691826.

Richani D, Dunning KR, Thompson JG, Gilchrist RB. Metabolic co-dependence of the oocyte and cumulus cells: essential role in determining oocyte developmental competence. Hum Reprod Update 2021;27: 27–47.

Richani D, Gilchrist RB. Approaches to oocyte meiotic arrest *in vitro* and impact on oocyte developmental competence. Biology of Reproduction 2021.

Richani D, Lavea CF, Kanakkaparambil R, Riepsamen AH, Bertoldo MJ, Bustamante S, Gilchrist RB. Participation of the adenosine salvage pathway and cyclic AMP modulation in oocyte energy metabolism. Scientific reports 2019;9: 18395.

Richani D, Poljak A, Wang B, Mahbub SB, Biazik J, Campbell JM, Habibalahi A, Stocker WA, Marinova MB, Nixon B et al. Oocyte and cumulus cell cooperativity and metabolic plasticity under the direction of oocyte paracrine factors. Am J Physiol Endocrinol Metab 2024;326: E366–e381.

Rodríguez-Nuevo A, Torres-Sanchez A, Duran JM, De Guirior C, Martínez-Zamora MA, Böke E. Oocytes maintain ROS-free mitochondrial metabolism by suppressing complex I. Nature 2022;607: 756–761.

Sanchez F, Le AH, Ho VNA, Romero S, Van Ranst H, De Vos M, Gilchrist RB, Ho TM, Vuong LN, Smitz J. Biphasic *in vitro* maturation (CAPA-IVM) specifically improves the developmental capacity of oocytes from small antral follicles. Journal of assisted reproduction and genetics 2019;36: 2135–2144.

Sánchez F, Lolicato F, Romero S, De Vos M, Van Ranst H, Verheyen G, Anckaert E, Smitz JEJ. An improved IVM method for cumulus-oocyte complexes from small follicles in polycystic ovary syndrome patients enhances oocyte competence and embryo yield. *Human reproduction (Oxford*, England) 2017;32: 2056–2068.

Sanfins A, Plancha CE, Albertini DF. Pre-implantation developmental potential from *in vivo* and *in vitro* matured mouse oocytes: a cytoskeletal perspective on oocyte quality. Journal of assisted reproduction and genetics 2015;32: 127–136.

Schouten N, Wang R, Torrance H, Van Tilborg T, Bastu E, Bergh C, D’Hooghe T, Friis Petersen J, Jayaprakasan K, Khalaf Y et al. Development and validation of a gonadotropin dose selection model for optimized ovarian stimulation in IVF/ICSI: an individual participant data meta-analysis. Hum Reprod Update 2025;31: 116–132.

Sciorio R, Miranian D, Smith GD. Non-invasive oocyte quality assessment. Biology of Reproduction 2022;106: 274–290.

Sha QQ, Zhu YZ, Li S, Jiang Y, Chen L, Sun XH, Shen L, Ou XH, Fan HY. Characterization of zygotic genome activation-dependent maternal mRNA clearance in mouse. Nucleic acids research 2020;48: 879–894.

Stocker WA, Walton KL, Richani D, Chan KL, Beilby KH, Finger BJ, Green MP, Gilchrist RB, Harrison CA. A variant of human growth differentiation factor-9 that improves oocyte developmental competence. J Biol Chem 2020;295: 7981–7991.

Su YQ, Sugiura K, Woo Y, Wigglesworth K, Kamdar S, Affourtit J, Eppig JJ. Selective degradation of transcripts during meiotic maturation of mouse oocytes. Developmental biology 2007;302: 104–117.

Sun H, Sun G, Zhang H, An H, Guo Y, Ge J, Han L, Zhu S, Tang S, Li C et al. Proteomic Profiling Reveals the Molecular Control of Oocyte Maturation. Mol Cell Proteomics 2023;22: 100481.

Tan TCY, Brown HM, Thompson JG, Mustafa S, Dunning KR. Optical imaging detects metabolic signatures associated with oocyte quality†. Biol Reprod 2022;107: 1014–1025.

Tan TCY, Dunning KR. Non-invasive assessment of oocyte developmental competence. Reproduction, fertility, and development 2022;35: 39–50.

Trapphoff T, Heiligentag M, Simon J, Staubach N, Seidel T, Otte K, Fröhlich T, Arnold GJ, Eichenlaub-Ritter U. Improved cryotolerance and developmental potential of *in vitro* and *in vivo* matured mouse oocytes by supplementing with a glutathione donor prior to vitrification. Molecular human reproduction 2016;22: 867–881.

Vigone G, Merico V, Prigione A, Mulas F, Sacchi L, Gabetta M, Bellazzi R, Redi CA, Mazzini G, Adjaye J et al. Transcriptome based identification of mouse cumulus cell markers that predict the developmental competence of their enclosed antral oocytes. BMC Genomics 2013;14: 380.

Vuong LN, Ho VNA, Ho TM, Dang VQ, Phung TH, Giang NH, Le AH, Pham TD, Wang R, Smitz J et al. In-vitro maturation of oocytes versus conventional IVF in women with infertility and a high antral follicle count: a randomized non-inferiority controlled trial. Human reproduction (Oxford, England) 2020a;35: 2537–2547.

Vuong LN, Ho VNA, Le AH, Nguyen NT, Pham TD, Nguyen MHN, Le HL, Le TK, Ha AN, Le XTH et al. Hormone-free vs. follicle-stimulating hormone–primed infertility treatment of women with polycystic ovary syndrome using biphasic *in vitro* maturation: a randomized controlled trial. Fertil Steril 2025;123: 253–261.

Vuong LN, Le AH, Ho VNA, Pham TD, Sanchez F, Romero S, De Vos M, Ho TM, Gilchrist RB, Smitz J. Live births after oocyte *in vitro* maturation with a prematuration step in women with polycystic ovary syndrome. Journal of assisted reproduction and genetics 2020b;37: 347–357.

Walls ML, Hunter T, Ryan JP, Keelan JA, Nathan E, Hart RJ. *In vitro* maturation as an alternative to standard *in vitro* fertilization for patients diagnosed with polycystic ovaries: a comparative analysis of fresh, frozen and cumulative cycle outcomes. Human Reproduction 2014;30: 88–96.

Wang Y, Zhang Y, Li T, Ren Y, Zhou P, Fu L, Xiao C, Huang Z, Huang H, Xie W et al. Transcriptional insights on the incomplete cytoplasmic maturation and developmental potential of oocytes cultured without granulosa cells in mice. BMC Genomics 2025;26: 270.

Wasserzug Pash P, Karavani G, Reich E, Zecharyahu L, Kay Z, Bauman D, Mordechai-Daniel T, Imbar T, Klutstein M. Pre-pubertal oocytes harbor altered histone modifications and chromatin configuration. Frontiers in cell and developmental biology 2022;10: 1060440.

Williams JPC, Auchynnikava T, McCarthy A, Ojarikre OA, Teresa Bertran M, Weston AE, Leonce D, Olsen J, Niakan KK, Skehel M et al. Dissecting PADI6 function defines oocyte cytoplasmic lattices as regulatory hubs for fundamental cellular processes. bioRxiv 2025: 2025.2002.2021.639491.

Xing J, Qiao G, Luo X, Liu S, Chen S, Ye G, Zhang C, Yi J. Ferredoxin 1 regulates granulosa cell apoptosis and autophagy in polycystic ovary syndrome. Clin Sci (Lond) 2023;137: 453–468.

Zaffagnini G, Cheng S, Salzer MC, Pernaute B, Duran JM, Irimia M, Schuh M, Böke E. Mouse oocytes sequester aggregated proteins in degradative super-organelles. Cell 2024;187: 1109–1126.e1121.

Zaffagnini G, Solé M, Duran JM, Polyzos NP, Böke E. The proteostatic landscape of healthy human oocytes. The EMBO journal 2025;44: 4611–4630.

Zhang H, Ji S, Zhang K, Chen Y, Ming J, Kong F, Wang L, Wang S, Zou Z, Xiong Z et al. Stable maternal proteins underlie distinct transcriptome, translatome, and proteome reprogramming during mouse oocyte-to-embryo transition. Genome Biol 2023;24: 166.

Zhang Y, Yan Z, Qin Q, Nisenblat V, Chang HM, Yu Y, Wang T, Lu C, Yang M, Yang S et al. Transcriptome Landscape of Human Folliculogenesis Reveals Oocyte and Granulosa Cell Interactions. Molecular cell 2018;72: 1021–1034.e1024.

Zhao Y, Liao X, Krysta AE, Bertoldo MJ, Richani D, Gilchrist RB. Capacitation IVM improves cumulus function and oocyte quality in minimally stimulated mice. Journal of assisted reproduction and genetics 2020;37: 77–88.

Zhu S, Li J, Wang X, Jin Y, Wang H, An H, Sun H, Han L, Shen B, Wang Q. The chromatin accessibility landscape of mouse oocytes during configuration transition. Cell Prolif 2025;58: e13733.

