## Supplemental Materials for "A proteomic signature of oocyte quality from models of varying oocyte developmental competence"

| Number of cumulus-oocyte complexes (COCs) per replicate | | | |
| --- | --- | --- | --- |
|  | In Vivo | CAPA-IVM | Trad-IVM |
| Replicate 1 | 277 | 198 | 198 |
| Replicate 2 | 214 | 156 | 150 |
| Replicate 3 | 200 | 106 | 106 |
| Replicate 4 | 316 | 126 | 123 |

*Supplementary Table 1. Number of cumulus-oocyte complexes (COCs) per replicate used for label-free proteomics.*

| **Antibody** | **Source/Reference** | **Catalogue no.** | **RRID** | **Dilution (Concentration)** |
| --- | --- | --- | --- | --- |
| Anti-ADX (FDX1) Rabbit Monoclonal | Abcam | ab108257 | AB_10862209 | 1:100 (9.28ug/mL) |
| Anti-CLTB Rabbit Polyclonal | ThermoFisher Scientific | PA5-109557 | AB_2854968 | 1:200 (4.55ug/mL) |
| Anti-EIF2A Rabbit Monoclonal | Abcam | ab169528 | AB_2819002 | 1:100 (4.16ug/mL) |
| Anti-PHGDH Rabbit Polyclonal | ThermoFisher Scientific | PA5-27578 | AB_2545054 | 1:500 (1.74ug/mL) |
| Anti-RPL24 Rabbit Polyclonal | ThermoFisher Scientific | PA5-30157 | AB_2547631 | 1:100 (10.20ug/mL) |

*Supplementary Table 2. Antibodies used in this study.*

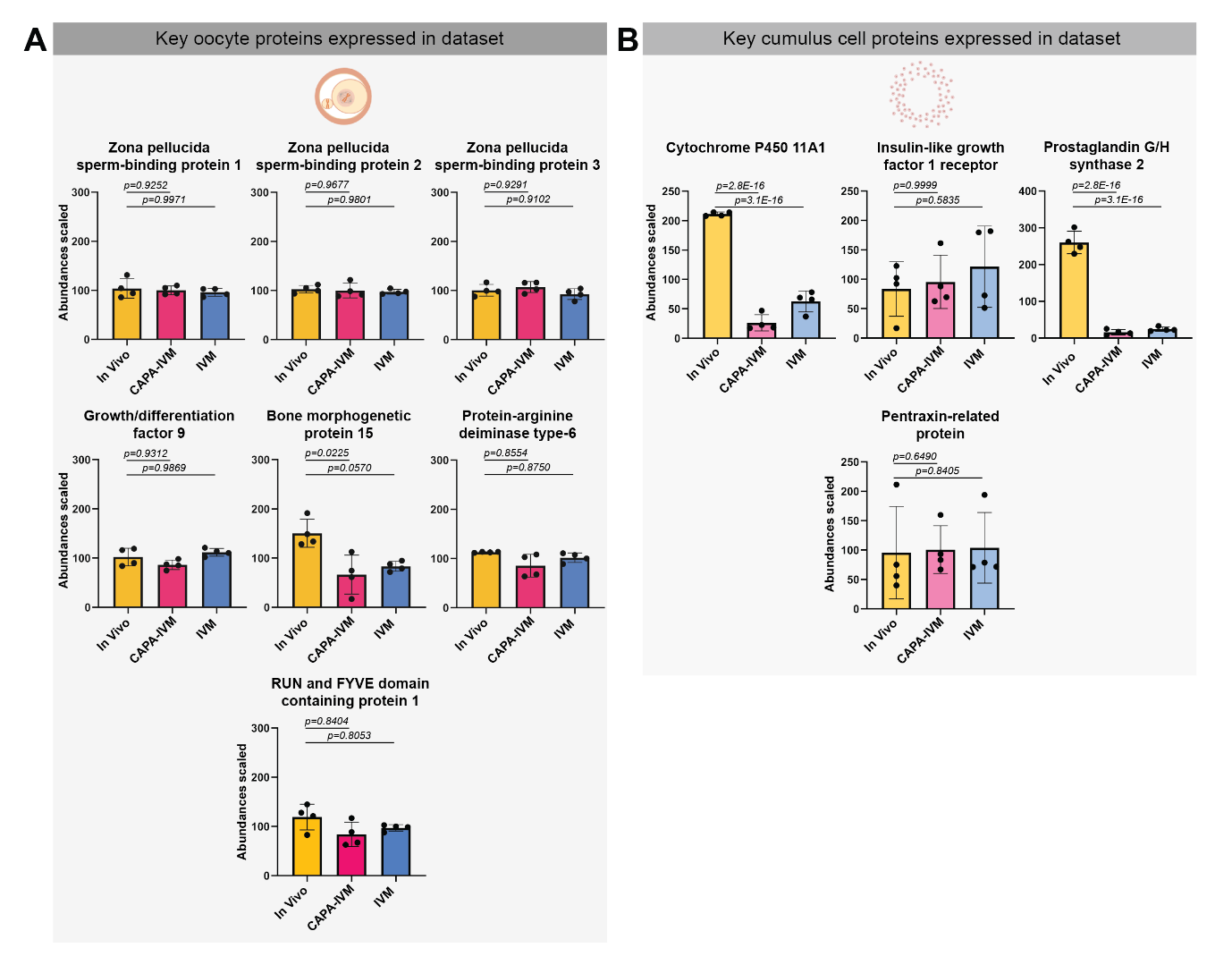

*Supplemental Figure 1. Key oocyte and cumulus cell protein expression across models of developmental competence.* Scaled proteomic abundances for individual proteins across *in vivo* (yellow), CAPA-IVM (pink) and IVM (blue) treatment groups expressed in A) oocytes and B) cumulus cells (n = 4, abundance ratio adjusted p-values and mean ± standard deviation).

**Datasets:**

**Supplemental Dataset 1**

Full list of proteins identified in oocytes by mass spectrometry and exported from Proteome Discoverer 2.4 software.

**Supplemental Dataset 2**

Full list of proteins identified in cumulus cells by mass spectrometry and exported from Proteome Discoverer 2.4 software.

**Supplemental Dataset 3**

Differential protein expression analysis in oocytes.

**Supplemental Dataset 4**

Differential protein expression analysis in cumulus cells.

**Supplemental Dataset 5**

List of proteins identified by pattern-based analysis in oocytes.

**Supplemental Dataset 6**

List of proteins identified by pattern-based analysis in cumulus cells.
